# Identification of 20 novel loci associated to ischemic stroke. Epigenome-Wide Association Study

**DOI:** 10.1101/2019.12.11.872945

**Authors:** Carolina Soriano-Tárraga, Uxue Lazcano, Eva Giralt-Steinhauer, Carla Avellaneda-Gómez, Ángel Ois, Ana Rodríguez-Campello, Elisa Cuadrado-Godia, Alejandra Gomez-Gonzalez, Alba Fernández-Sanlés, Roberto Elosua, Israel Fernández-Cadenas, Natalia Cullell, Joan Montaner, Sebastian Moran, Manel Esteller, Jordi Jiménez-Conde, Jaume Roquer

## Abstract

**Rationale:** DNA methylation is dynamic, varies throughout the life course, and its levels are influenced by lifestyle and environmental factors, as well as by genetic variation. The leading genetic variants at stroke risk loci identified to date explain roughly 1–2% of stroke heritability. Most of these single nucleotide polymorphisms are situated within a regulatory sequence marked by DNase I hypersensitivity sites, which would indicate involvement of an epigenetic mechanism.

**Objective:** To detect epigenetic variants associated to stroke occurrence and stroke subtypes.

**Methods and Results:** A two-stage case-control epigenome-wide association study was designed. The discovery sample with 401 samples included 218 ischemic stroke (IS) patients, assessed at Hospital del Mar (Barcelona, Spain) and 183 controls from the REGICOR cohort. In two independent samples (N=226 and N=166), we replicated 22 CpG sites differentially methylated in IS in 21 loci, including 2 CpGs in locus *ZFHX3*, which includes known genetic variants associated with stroke. The pathways associated to these loci are inflammation and angiogenesis. The meta-analysis identified 384 differentially methylated CpGs, including loci of known stroke and vascular risk genetic variants, enriched by loci involved in lipid metabolism, adipogenesis, circadian clock, and glycolysis pathways. Stratification analysis by stroke subtypes revealed distinct methylation patterns.

**Conclusions:** We identified a set of 22 CpGs in 21 loci associated with IS. Our analysis suggests that DNA methylation changes may contribute to orchestrating gene expression that contributes to IS.

## INTRODUCTION

Ischemic stroke (IS), the etiology of 80% of all strokes, is a complex and heterogeneous disease with high rates of mortality and long-term disability. Stroke pathogenesis involves a number of different disease processes as well as interactions between environmental, vascular, systemic, genetic, and central nervous system factors ^1^.

In recent years, genome-wide association studies (GWAS) have identified many genes robustly associated to IS and its subtypes which show a distinct genetic background, but the underlying genes and pathways are not yet fully understood. Half of the identified genes share a genetic association with other vascular traits; the greatest correlation is with blood pressure. However, the underlying biological pathways do not seem to involve known vascular risk factors and thus the new pathways may constitute new targets for stroke prevention ^2^.

The heritability of stroke, as calculated from genome-wide data, has been estimated to be 30–40% ^3–5^. The lead variants at stroke risk loci identified to date explain roughly 1–2% of this heritability and their number remains relatively small compared to other common conditions, such as coronary artery disease ^6^. Moreover, most of these leading single nucleotide polymorphisms (SNPs) reside within intergenic or intronic regions, and most are situated within a regulatory sequence marked by DNase I hypersensitivity sites, which would indicate the involvement of an epigenetic mechanism ^2,7,8^.

DNA methylation (DNAm), an epigenetic mechanism essential for regulation of gene expression, consists of the covalent addition of a methyl group to a cytosine nucleotide, primarily in the context of a CpG dinucleotide. DNAm is dynamic, varies throughout the life course, and its levels are influenced by lifestyle and environmental factors, as well as by genetic variation ^9^.

Stroke epigenetics research has revealed that stroke patients are globally hypomethylated and are *epigenetically* older than healthy controls. This accelerated aging is strongly associated with 3-month IS outcome and mortality, independently of health profile, lifestyle, and genetic characteristics ^10–13^.

We hypothesized that DNA methylation markers would identify additional stroke risk loci and improve the mapping of genomic variability, providing new insights into stroke pathology. The aim of the present study was to detect epigenetic variants associated to IS occurrence and stroke subtypes.

## METHODS

The data that support the findings of this study are available from the corresponding author upon reasonable request.

### Study design

A two-stage case-control epigenome-wide association study (EWAS) was designed, including discovery and replication analyses. The discovery sample consisted of 401 samples (183 controls and 218 IS patients). Two independent samples (N=392,226 from Hospital del Mar [MAR-2] and 166 from Hospital Vall d’Hebron [HVH]) were used to replicate the top 500 methylation-variable positions (MVPs) identified in the discovery sample with an arbitrary p-value ≤7.1×10^−6^. Joint metaanalysis of all 3 samples was performed (N=793). Stratification analysis by Trial of Org 10172 in Acute Stroke Treatment (TOAST) was performed. Functional and pathway analyses were conducted.

### Study Sample

European ancestry controls and patients with a diagnosis of IS according to World Health Organization criteria were selected.

### Epigenome-Wide Association Studies

After EWAS quality controls, association analysis was performed in controls and IS samples using a multivariate linear regression. The top MVPs identified in the discovery stage were analyzed in the validation samples. The results were joined in a weighted z-score meta-analysis. Stratification analysis by TOAST criteria was performed using a multivariate linear regression for large-artery atherosclerosis (LAA), cardioembolic (CE), small vessel disease (SVD), and undetermined (UND) etiologies. Functional and pathway analyses were conducted. All MVPs with p-values <1.39×10^−7^ were considered statistically significant for a genome-wide approach.

## RESULTS

### Discovery stage. Genome-wide effect of ischemic stroke on methylation status

Of the 865,859 initial CpG sites in Infinium MethylationEPIC Beadchip, a total of 358,709 (41.4%) that passed the quality controls and were common in HumanMethylation450 Beadchip were included in the discovery analysis (**Table S1-2**). Genome-wide DNAm analysis from whole blood was performed in the discovery sample (N=401). Clinical and demographic differences between the IS and control individuals are shown in **Table 1**.

**Table 1.**
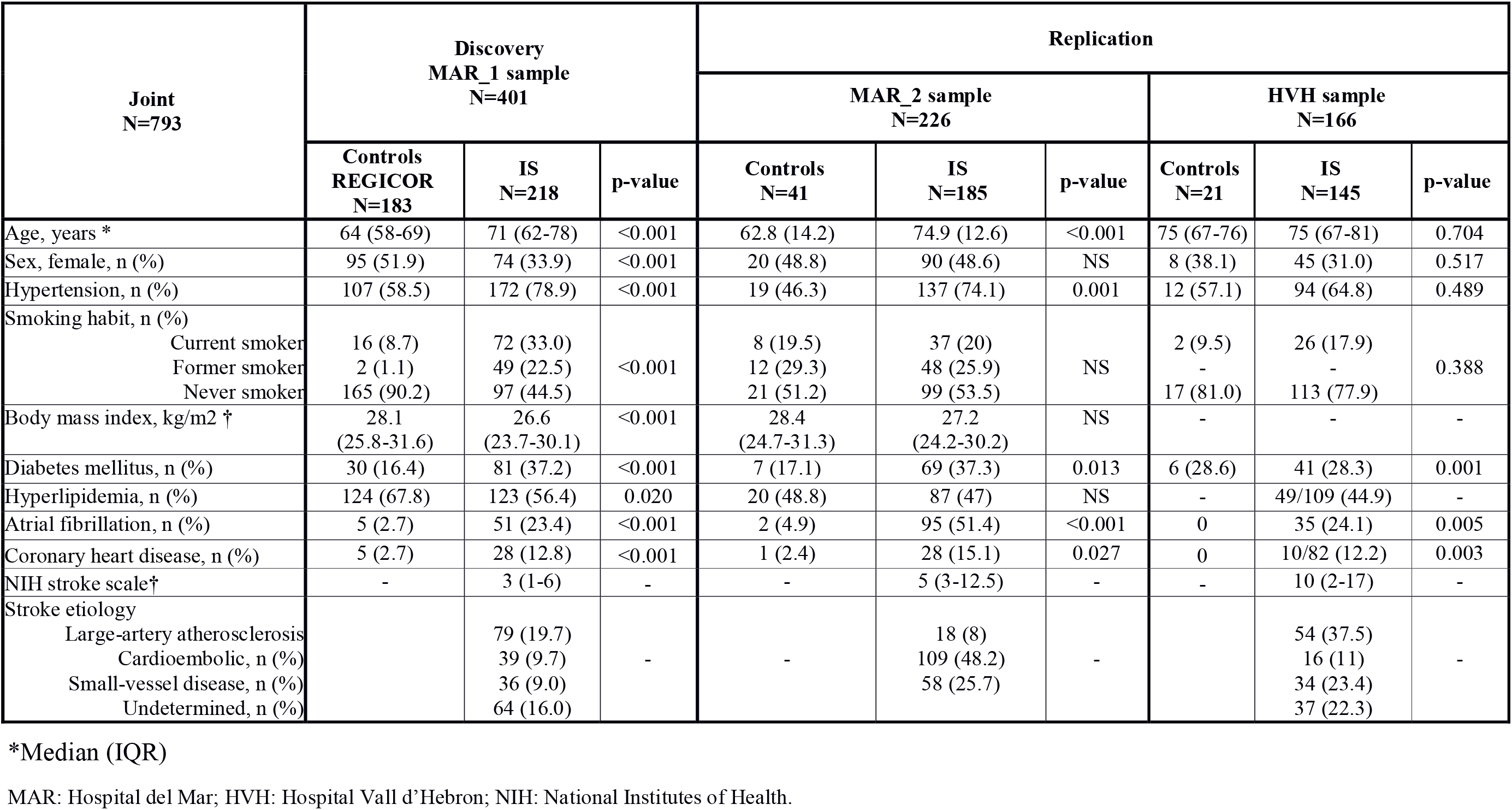
Descriptive characteristics of the samples.

| Joint<br>N=793 | Discovery<br>MAR_1 sample<br>N=401 |  |  | Replication |  |  |  |  |  |
| --- | --- | --- | --- | --- | --- | --- | --- | --- | --- |
|  |  |  |  | MAR_2 sample<br>N=226 |  |  | HVH sample<br>N=166 |  |  |
|  | Controls<br>REGICOR<br>N=183 | IS<br>N=218 | p-value | Controls<br>N=41 | IS<br>N=185 | p-value | Controls<br>N=21 | IS<br>N=145 | p-value |
| Age, years * | 64 (58-69) | 71 (62-78) | <0.001 | 62.8 (14.2) | 74.9 (12.6) | <0.001 | 75 (67-76) | 75 (67-81) | 0.704 |
| Sex, female, n (%) | 95 (51.9) | 74 (33.9) | <0.001 | 20 (48.8) | 90 (48.6) | NS | 8 (38.1) | 45 (31.0) | 0.517 |
| Hypertension, n (%) | 107 (58.5) | 172 (78.9) | <0.001 | 19 (46.3) | 137 (74.1) | 0.001 | 12 (57.1) | 94 (64.8) | 0.489 |
| Smoking habit, n (%) |  |  |  |  |  |  |  |  |  |
| Current smoker | 16 (8.7) | 72 (33.0) | <0.001 | 8 (19.5) | 37 (20) | NS | 2 (9.5) | 26 (17.9) | 0.388 |
| Former smoker | 2 (1.1) | 49 (22.5) |  | 12 (29.3) | 48 (25.9) |  | - | - |  |
| Never smoker | 165 (90.2) | 97 (44.5) |  | 21 (51.2) | 99 (53.5) |  | 17 (81.0) | 113 (77.9) |  |
| Body mass index, kg/m2 † | 28.1<br>(25.8-31.6) | 26.6<br>(23.7-30.1) | <0.001 | 28.4<br>(24.7-31.3) | 27.2<br>(24.2-30.2) | NS | - | - | - |
| Diabetes mellitus, n (%) | 30 (16.4) | 81 (37.2) | <0.001 | 7 (17.1) | 69 (37.3) | 0.013 | 6 (28.6) | 41 (28.3) | 0.001 |
| Hyperlipidemia, n (%) | 124 (67.8) | 123 (56.4) | 0.020 | 20 (48.8) | 87 (47) | NS | - | 49/109 (44.9) | - |
| Atrial fibrillation, n (%) | 5 (2.7) | 51 (23.4) | <0.001 | 2 (4.9) | 95 (51.4) | <0.001 | 0 | 35 (24.1) | 0.005 |
| Coronary heart disease, n (%) | 5 (2.7) | 28 (12.8) | <0.001 | 1 (2.4) | 28 (15.1) | 0.027 | 0 | 10/82 (12.2) | 0.003 |
| NIH stroke scale† | - | 3 (1-6) | - | - | 5 (3-12.5) | - | - | 10 (2-17) | - |
| Stroke etiology |  |  |  |  |  |  |  |  |  |
| Large-artery atherosclerosis |  | 79 (19.7) | - | - | 18 (8) | - |  | 54 (37.5) | - |
| Cardioembolic, n (%) |  | 39 (9.7) |  |  | 109 (48.2) |  |  | 16 (11) |  |
| Small-vessel disease, n (%) |  | 36 (9.0) |  |  | 58 (25.7) |  |  | 34 (23.4) |  |
| Undetermined, n (%) |  | 64 (16.0) |  |  |  |  |  | 37 (22.3) |  |

\*Median (IQR)
MAR: Hospital del Mar; HVH: Hospital Vall d'Hebron; NIH: National Institutes of Health.

Top 500 CpG sites showing a suggestive association with methylation differences between IS and control samples were selected for replication, with an arbitrary p-value ≤7.1×10^−6^ (**Supplementary Table S3**). Manhattan and QQ plots are shown in **Figure 1**.

**Figure 1.**
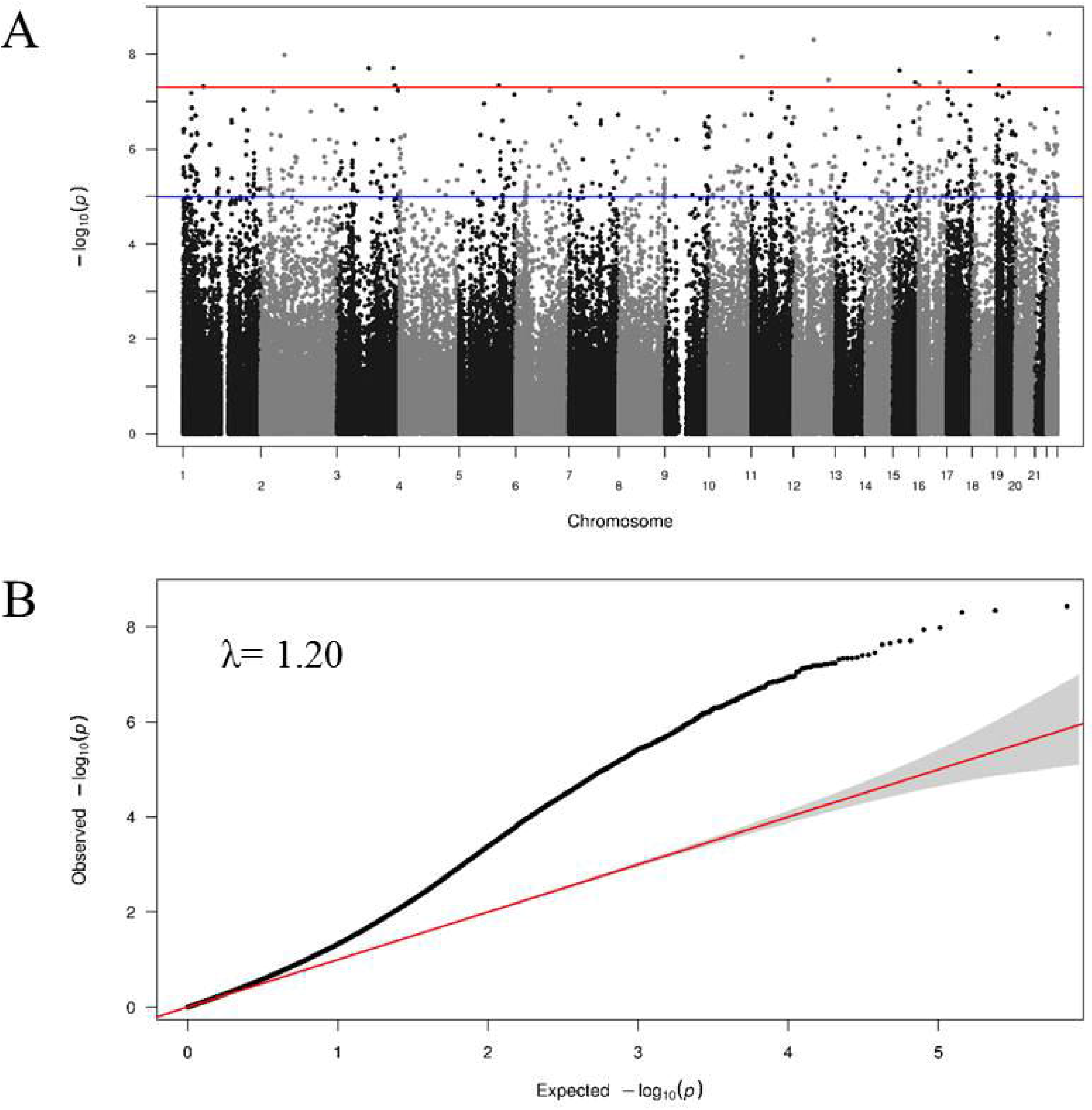
Manhattan and QQ-plot in the discovery sample. **A)** Manhattan plot (red line and blue line for statistically and nominally significance threshold) and B) QQ-plot.

### Replication stage

After applying the same quality control steps and normalization as in the discovery analysis, the validation analysis included 392 individuals (226 from MAR-2 and 166 from HVH). The characteristics of the populations included in the replication stage are shown in **Table 1-2**. We replicated 93 CpGs in one replication sample and 114 CpGs in the other; 22 MVPs in 21 loci were replicated in both samples. All of the 22 MVPs were hypomethylated in all the samples, except the one located at *MAPK1;* however, IS samples showed significantly higher methylation levels, compared to controls (**Table 2, Supplementary Figure S1)**. Function and traits associated to the 21 loci that harbor the 22 CpGs identified are described in **Supplementary Table S4-7.**

**Table 2.** CpG sites differentially methylated between ischemic stroke and controls in discovery sample validated in two replication samples and the meta-analysis.

| Features |  |  |  |  | Discovery |  |  |  |  | Replication 1 |  | Replication 2 |  | Meta-analysis |  |
| --- | --- | --- | --- | --- | --- | --- | --- | --- | --- | --- | --- | --- | --- | --- | --- |
| CpG | Chr | Pos (hg37) | CpG location | Gene | LogFC | Beta (SD) Control | Beta (SD) IS | DNAm diff | P-value | LogFC | P-value FDR | LogFC | P-value FDR | P-value | Dir |
| cg09915769 | 19 | 7660977 | Island | CAMSAP3 | 0.03 | 0.07 (0.02) | 0.09 (0.02) | 3 | 4.56E-08 | 4.56E-08 | 5.08E-05 | 0.02 | 5.59E-06 | 4.26E-16 | +++ |
| cg02463426 | 19 | 16683387 | Island | SLC35E1 | 0.02 | 0.06 (0.04) | 0.10 (0.06) | 2 | 3.11E-07 | 3.11E-07 | 1.34E-05 | 0.02 | 2.85E-06 | 5.19E-16 | +++ |
| cg00614832 | 16 | 73092394 | Island | ZFHX3 | 0.04 | 0.07 (0.04) | 0.11 (0.07) | 4 | 6.41E-07 | 6.41E-07 | 0.0001 | 0.04 | 2.15E-08 | 3.23E-16 | +++ |
| cg23962478 | 22 | 50354086 | Island | PIM3 | 0.03 | 0.06 (0.03) | 0.08 (0.04) | 3 | 8.31E-07 | 8.31E-07 | 9.94E-05 | 0.02 | 1.14E-06 | 5.61E-15 | +++ |
| cg23681311 | 22 | 22221878 | Island | MAPK1 | 0.04 | 0.78 (0.03) | 0.77 (0.03) | 4 | 8.50E-07 | 8.50E-07 | 0.0001 | 0.05 | 2.18E-05 | 6.59E-14 | +++ |
| cg13696351 | 9 | 140063617 | Island | LRRC26 | 0.03 | 0.05 (0.03) | 0.08 (0.04) | 3 | 9.07E-07 | 9.07E-07 | 3.60E-06 | 0.03 | 1.45E-06 | 3.03E-16 | +++ |
| cg01182555 | 14 | 62162064 | Island | HIF1A | 0.02 | 0.10 (0.03) | 0.13 (0.04) | 2 | 9.57E-07 | 9.57E-07 | 0.0001 | 0.004 | 2.73E-05 | 8.18E-14 | +++ |
| cg07691609 | 19 | 662740 | Island | RNF126 | 0.02 | 0.06 (0.04) | 0.12 (0.08) | 2 | 1.16E-06 | 1.16E-06 | 8.56E-05 | 0.02 | 6.88E-08 | 9.84E-16 | +++ |
| cg01733795 | 17 | 7465439 | Island | SENP3 | 0.02 | 0.10 (0.04) | 0.13 (0.06) | 2 | 1.38E-06 | 1.38E-06 | 2.41E-06 | 0.02 | 1.19E-06 | 2.96E-16 | +++ |
| cg08184047 | 17 | 79849980 | Island | ANAPC11 | 0.02 | 0.06 (0.04) | 0.08 (0.06) | 2 | 1.74E-06 | 1.74E-06 | 4.14E-07 | 0.01 | 1.00E-04 | 3.24E-15 | +++ |
| cg04355250 | 12 | 113796401 | Island | PLBD2 | 0.02 | 0.07 (0.02) | 0.08 (0.04) | 2 | 1.93E-06 | 1.93E-06 | 6.47E-05 | 0.01 | 1.33E-05 | 5.78E-14 | +++ |
| cg16573386 | 1 | 1334508 | Island | CCNL2 | 0.02 | 0.10 (0.03) | 0.12 (0.04) | 2 | 2.67E-06 | 2.67E-06 | 1.09E-06 | 0.02 | 4.31E-08 | 2.90E-17 | +++ |
| cg23281075 | 1 | 31538727 | Island | PUM1 | 0.02 | 0.08 (0.02) | 0.11 (0.05) | 2 | 2.84E-06 | 2.84E-06 | 9.06E-05 | 0.02 | 6.63E-07 | 1.38E-14 | +++ |
| cg07786668 | 16 | 73092391 | Island | ZFHX3 | 0.04 | 0.09 (0.03) | 0.11 (0.04) | 4 | 3.06E-06 | 3.06E-06 | 3.70E-08 | 0.06 | 3.57E-08 | 1.63E-18 | +++ |
| cg04482794 | 1 | 226925181 | Island | ITPKB | 0.02 | 0.07 (0.04) | 0.12 (0.07) | 2 | 3.36E-06 | 3.36E-06 | 2.77E-05 | 0.01 | 4.32E-06 | 2.05E-14 | +++ |
| cg07806715 | 19 | 48018254 | Island | NAPA | 0.02 | 0.05 (0.03) | 0.09 (0.05) | 2 | 3.58E-06 | 3.58E-06 | 1.49E-06 | 0.02 | 4.94E-07 | 3.14E-16 | +++ |
| cg08676905 | 10 | 6019609 | Island | IL15RA | 0.02 | 0.05 (0.02) | 0.07 (0.02) | 2 | 3.80E-06 | 3.80E-06 | 2.82E-06 | 0.02 | 1.69E-06 | 1.47E-15 | +++ |
| cg07619799 | 4 | 185747409 | Island | ACSL1 | 0.02 | 0.04 (0.02) | 0.07 (0.04) | 2 | 4.48E-06 | 4.48E-06 | 7.15E-05 | 0.01 | 1.53E-05 | 1.79E-13 | +++ |
| cg04759220 | 5 | 78532560 | Island | JMY | 0.03 | 0.07 (0.05) | 0.10 (0.07) | 2 | 4.70E-06 | 4.70E-06 | 5.26E-06 | 0.02 | 2.54E-05 | 2.66E-14 | +++ |
| cg01963056 | 8 | 144911482 | Island | PUF60 | 0.01 | 0.02 (0.01) | 0.03 (0.02) | 1 | 6.12E-06 | 6.12E-06 | 6.67E-06 | 0.07 | 2.45E-12 | 1.21E-13 | +++ |
| cg25869317 | 15 | 101792241 | Island | CHSY1 | 0.04 | 0.07 (0.02) | 0.08 (0.03) | 4 | 7.00E-06 | 7.00E-06 | 1.67E-06 | 0.04 | 1.72E-10 | 3.06E-19 | +++ |
| cg04192862 | 10 | 28966472 | Island | BAMBI | 0.03 | 0.07 (0.03) | 0.09 (0.04) | 3 | 7.10E-06 | 7.10E-06 | 2.19E-07 | 0.01 | 5.59E-06 | 6.32E-19 | +++ |

CpG, CpG identification; Chr, chromosome location; pos, genomic position; gene, associated gene and observed mean $\beta$ -values (standard deviation); LogFC, estimate of the log2-fold-change corresponding to the effect or contrast; P-value FDR, false discovery rate correcting by multiple testing; Dir, direction of the effect. All p-values were calculated with beta-value.

We used the GeneMANIA algorithm on the 21 candidate genes (*CAMSAP3, SLC35E1, ZFHX3, PIM3, MAPK1, LRRC26, HIF1A, RNF126, SENP3, ANAPC11, PLBD2, CCNL2, PUM1, ITPKB, NAPA, IL15RA, ACSL1, JMY, PUF60, CHSY1, BAMBI*) and known IS-related genes^2^. We found that our candidate genes have genetic interactions and are co-expressed with known IS genes (**Supplementary Figure S2**). *FGA* and *ZFHX3* are involved in the same pathway, as are *HIF1A* and *FURIN, HIF1A* and *MAPK1*. Shared protein domains exist between *HIF1A* and *TWIST1, LRRC26* and *LRCH1, ANACPC11* and *RNF126, ZFHX3* and *ZNF318*, and *CHSY1* and *ABO. CDK6* shares with *MAPK1* and *PIM3*. Finally, there is co-localization expression of *ITPKB* and *SH3PXD2A*; *NAPA* and *RGS7; NAPA, RGS7, ZFHX3*, and *MAPK1; ZFHX3* and *TBX3;* and *IL15RA, CASZ1*, and *FOXF2*.

The 22 validated CpGs include 2 CpGs in *ZFHX3* loci that harbor known stroke variants. The pathway analysis showed *HIF1A* and *MAPK1* are associated to angiogenesis (FDR p-values=0.014). *IL15RA* and *HIF1A* are involved in inflammatory and interferon gamma response (FDR p-values=0.034). Additionally, *NAPA* and *MAPK1* are involved in protein secretion functions (FDR p-values=0.024) (**Supplementary Table S5**). GTEX data indicate that 20 loci (the exception being *LRRC26*) are expressed in blood vessels and brain (**Supplementary Figure S3**).

These 22 MVPs are associated to CpG islands: 20 are in an enhancer-associated position, 2 are associated to H3K27AC (indicating active gene expression), and 21 are located in a DNase I hypersensitive site, functionally related to transcriptional activity (**Supplementary Table S7**).

### Meta-analysis

Meta-analysis results of all 3 samples (N=793) are shown in **Supplementary Table S8**. A total of 384 MVPs were significant after Bonferroni correction (multiple testing). However, 30 (7.8%) of these MVPs were only present in two samples.

The 384 significantly associated CpGs include 4 CpGs in *ZFHX3, SH2B3* and *WNK1* loci, which harbor known stroke variants. The traits and diseases associated to loci containing these CpGs are shown in **Figure 2.** The functional analysis of the significant loci showed enrichment for genes involved in adipogenesis (FDR p-value=4.13×10^−4^), triglyceride and lipid metabolism (FDR p-value=0.044), circadian clock (FDR p-value=0.044), and regulation of glycolysis pathway (FDR p-value=0.037) (**Supplementary Tables S9-11**).

**Figure 2.**
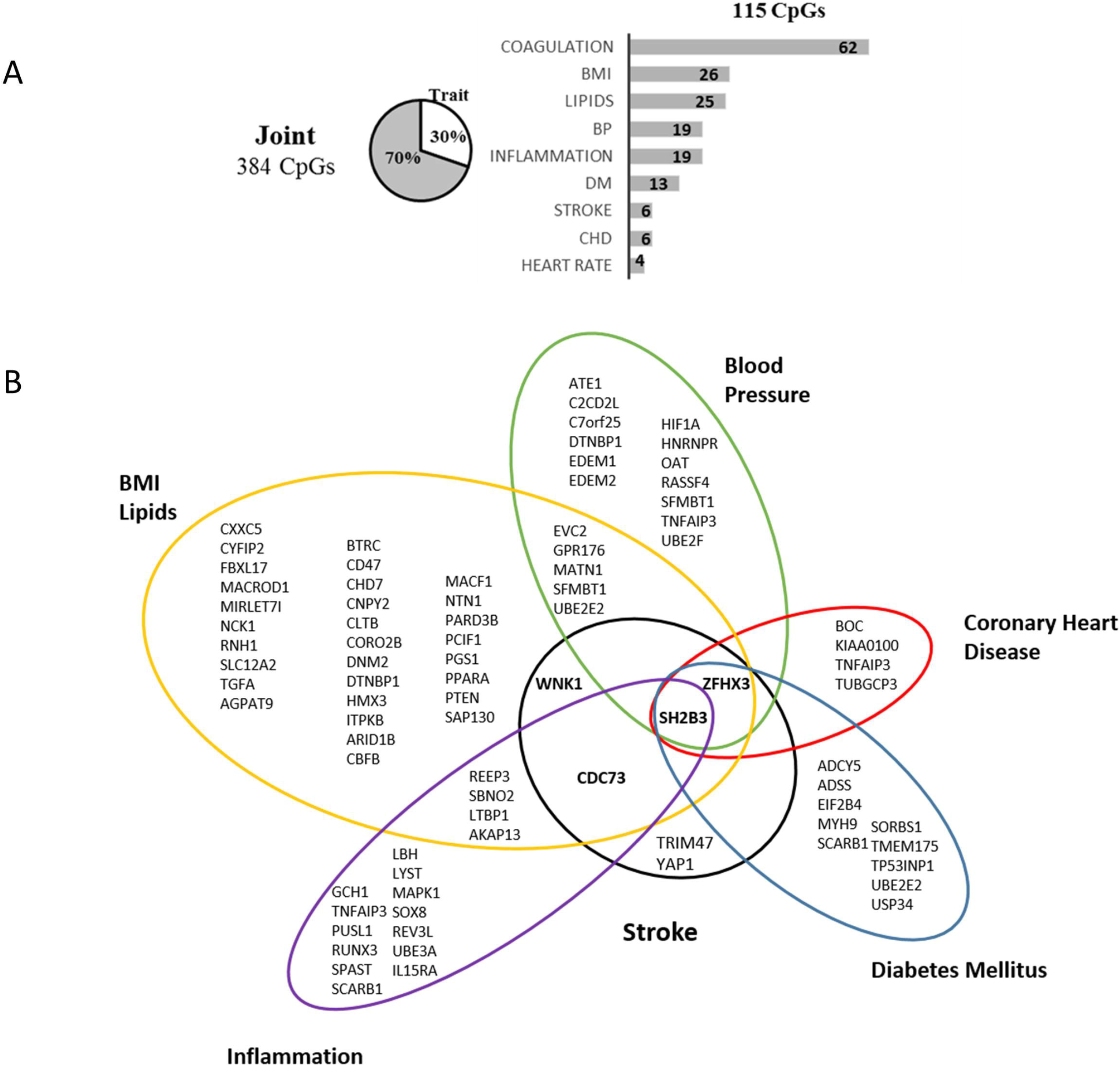
Summary of loci harboring significant CpGs in meta-analysis, overlapping known genome-wide associations to phenotypes and diseases. A) 115 CpG (30%) of the 384 joint CpGs are associated to known variants. Bars represent the number of loci associated to each trait; 37 loci are associated to more than 1 trait. B) List of loci with known associations. In bold, loci validated in discovery. BMI: Body Mass Index; BP: Blood Pressure; DM: Diabetes Mellitus; CHD: Coronary Heart Disease

### EWAS stratified analysis by IS subtypes

We combined the IS samples from the two Hospital del Mar samples (MAR_1 and MAR_2, N=627) to increase the statistical power to identify MVP sites associated to IS subtypes (TOAST classification), stratified by LAA, CE, SVD, and UND. Clinical and demographic characteristics and EWAS results are shown in **Supplementary Material (Table S12-16)**. Manhattan and QQ plots are in **Supplemental Figure S4**. In the meta-analysis, UND and SVD subtypes were the main contributors of significant IS-associated CpGs (**Figure 3**).

**Figure 3.**
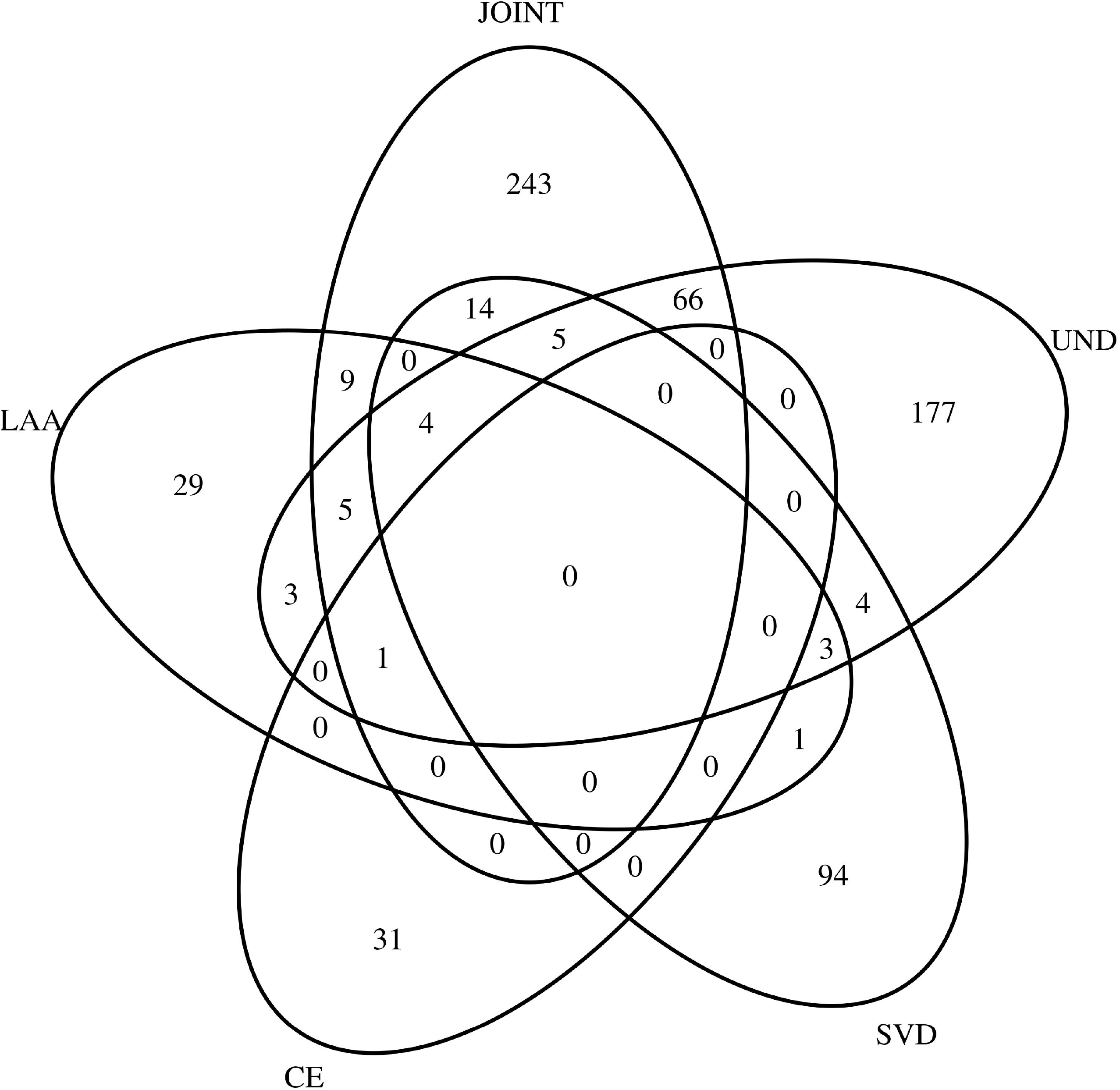
Venn Diagrams showing the overlap of methylation-variable positions between metaanalysis and the ischemic stroke subtypes. LAA, Large-Artery Atherosclerosis: CE, Cardiembolic; SVD, Small vessel diseases; UND, Undetermined IS subtypes.

In LAA subtype, we identified 57 MVPs suggestive of statistical significance (**Supplemental Table S13**). There was an enrichment for genes involved in plasma high-density lipoprotein cholesterol (HDL-C) levels (FDR p-values=0.039) and nominal significance for the adipogenesis pathway (p-values=0.0023) (**Supplementary Tables S17)**. In CE subtype, we identified 31 CpGs (**Supplemental Table S14**). The functional analysis showed the insulin signaling pathway as nominally significant (p-values=1.7×10^−3^) (**Supplementary Tables S18)**. In SVD subtype, we identified 121 MVPs suggestive of association (**Supplemental Table S15**). Enrichment analysis was significant for different traits associated to lipid metabolism (FDR p-values=0.013495) and nominally significant for adipogenous hallmark genes (p-value=0.0024) (**Supplemental Table S19**).

In UND subtype, we identified 294 MVPs (**Supplemental Table S16**). Enrichment analysis was significant for traits associated to inflammation (FDR p-values=0.0027) and cellular processes (**Supplementary Tables S20**).

The traits and diseases associated to loci that contain the suggestive CpGs associated to IS subtypes are shown in **Supplementary Tables S20-S24**. Approximately 30% of these MVPs overlap with known loci having genetic variants associated to vascular risk factors **(Figure 4)**.

**Figure 4.**
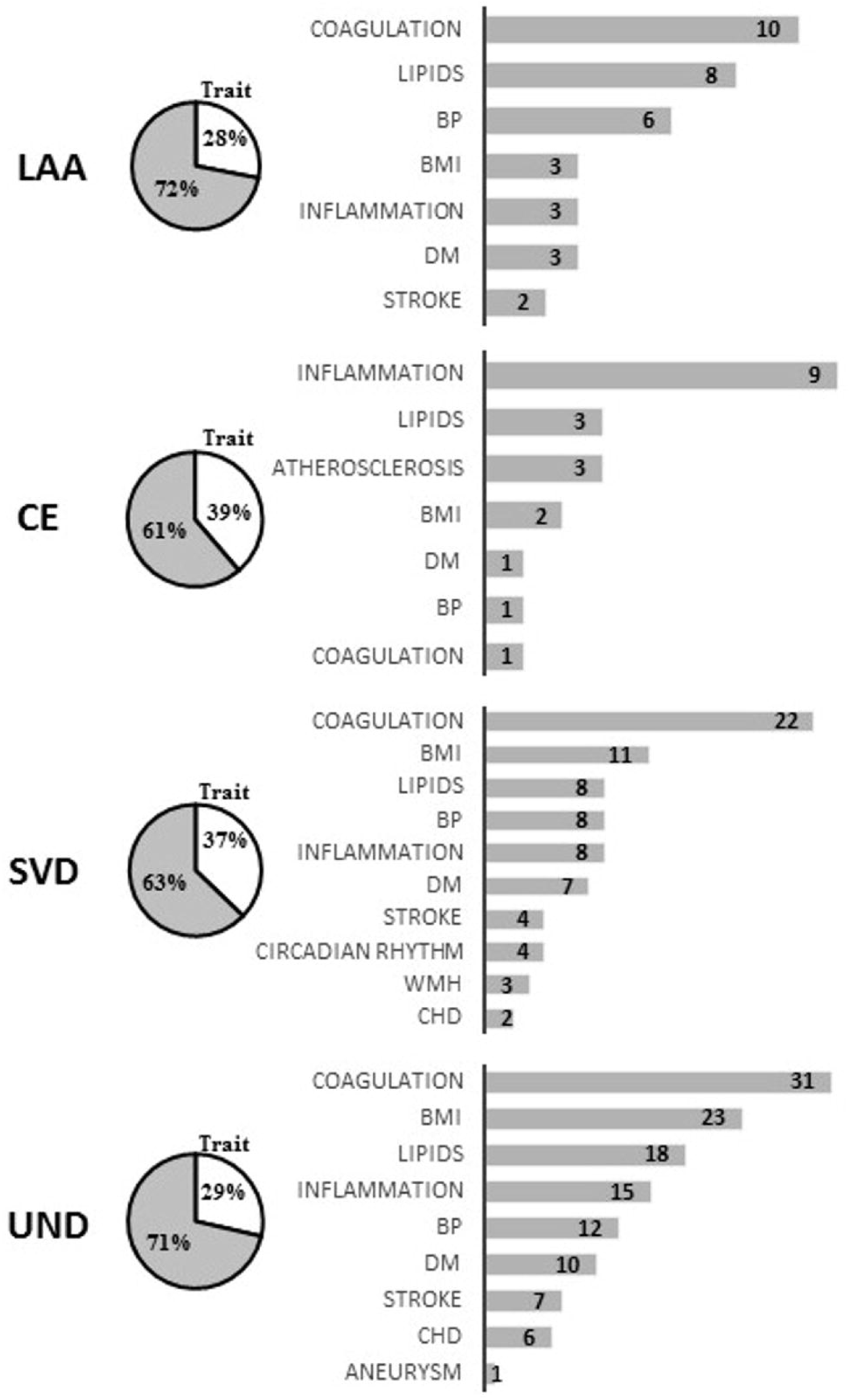
Summary of loci harboring nominally significant CpGs, by TOAST, associated to phenotypes and diseases in genome-wide association studies. One locus can be associated to more than one trait. LAA, Large-Artery Atherosclerosis: CE, Cardiembolic; SVD, Small vessel diseases; UND, Undetermined IS subtypes; BMI, Body Mass Index; BP, Blood Pressure; DM, Diabetes Mellitus; CHD, Coronary Heart Disease.

## Discussion

Novel findings in the present study include an association between IS and differentially methylated positions of 22 CpGs located in 21 loci (*CAMSAP3, SLC35E1, ZFHX3, PIM3, MAPK1, LRRC26, HIF1A, RNF126, SENP3, ANAPC11, PLBD2, CCNL2, PUM1, ITPKB, NAPA, IL15RA, ACSL1, JMY, PUF60, CHSY1, BAMBI)*. These CpGs are more methylated in IS than in control samples, except the one in the locus *MAPK1*. Two of these CpGs, cg00614832 and cg07786668, are situated in the *ZFHX3* locus, a known stroke gene^2^. Pathway analysis indicated that these 21 loci are involved in inflammation and angiogenesis pathways. Additionally, they share pathways, protein domains, genetic interaction and co-expression with stroke-associated genes identified in previous GWAS studies ^2^.

The repercussion of these findings in gene expression has not been studied; however, the 22 validated CpGs are located in CpG island-promoters, 20 of them in enhancer elements, and 21 in a DNase I hypersensitive site, functionally related to transcriptional activity; 12 are associated to H3K27AC, indicating active gene expression. All together, these results suggest a possible effect on gene expression.

The meta-analysis identified 384 MVPs significantly associated to IS, but replication is needed to confirm these results because of possible inflation of p-values. These MVPs were enriched for genes involved in adipogenesis, triglyceride and lipid metabolism, circadian clock, and regulation of glycolysis pathway. We also identified shared MVP associations with 5 genes (*SH2B3, ZFHX3, WNK1* (beside *NINJ2), PCIF1, TRIM47, YAP1*) previously associated to stroke, to vascular risk factors and to traits such as diabetes mellitus, coagulation, blood pressure, aortic aneurysm, body mass index, white-matter hyperintensity, and coronary heart disease^2,14–20^.

The overlap between loci containing SNPs associated to vascular risk factors and those related to stroke highlights the importance of epigenetics in identifying possible new biomarkers of stroke risk or/and new causal variants related to stroke. The question of potential independent effects of MVPs and SNPs remains to be studied. Most of the stroke risk SNPs identified are located in intergenic or intronic regions, and MVPs that were differentially methylated in the meta-analysis are located in the promoter region, making them better candidates to be the causal variant^2^.

The stratification analysis by stroke subtypes revealed distinct methylation patterns that could provide further mechanistic insights; however, replication is needed. The LAA subtype showed enrichment for genes involved in plasma HDL-C levels and adipogenesis pathway which correlates to LAA pathophysiology. Its top MVP, in the *GPS2* locus, has recently been associated with obesity and diabetes mellitus, with the authors suggesting that this gene may contribute to atherogenesis^21^. Additionally, cg17218495 is in the *SMARCA4* locus, previously associated to stroke and lipid metabolism)^22,23^. The CE analysis was enriched by genes involved in the insulin signaling pathway and adipogenous hallmark genes, while SVD analysis was enriched by genes involved in lipid metabolisms. Moreover, two MVPs were located in *CNNM2*, a known variant associated to blood pressure, and *TRIM47*, associated to white-matter hyperintensities which correlates to SVD pathophysiology ^24,25^.

DNAm can respond to changes in the environment and might mediate gene expression and phenotypes induced by prior exposure, such as glycemic exposure, a phenomenon called metabolic memory^26–28^. Based on those studies, DNAm may be one of the mechanisms involved in the metabolic memory. We only reported an association with IS traits, but cannot infer causality of these methylation changes; however, these MVPs could be causal variants, biomarkers of exposure, or stochastic changes. Further investigation is needed.

In this study, we analyzed one of the largest available series of IS patients with genome-wide methylation data. We replicated our results in two independent samples. Moreover, we identified 22 CpG sites related to IS, two of them located in the *ZFHX3* locus, harboring known genetic variants associated with stroke. Stratification by stroke subtype revealed distinct methylation patterns associated to different vascular risk factors; however, these results must be replicated.

EWAS are prone to significant inflation and bias of test statistics. Neither GWAS-based methodology nor confounder adjustment methods completely remove bias and inflation^29^. Epigenome data differ from genetic data: they are quantitative measures (whereas genotypes are discrete) that are subject to major confounding effects of batch analysis, whether technical, such as DNA extraction protocols; biological, including cellular heterogeneity; or epigenetic changes caused by the disease process itself ^29–32^. In our study, blood samples were collected in the acute phase of IS; although we adjusted for stroke severity using the NIH scale in an effort to correct the residual confounding, the inflation is evident. Nonetheless, some signals were identified and others require replication.

Some limitations of the study should be considered. First, we measured DNAm in peripheral blood cells. Some authors have suggested that the methylation levels of some CpGs/regions are tissuespecific ^33^, and we might have lost some signals by not choosing specific tissues that could have a higher impact in DNAm. However, other authors consider methylation patterns of whole blood a good proxy for the methylation levels from a specific site of action. Second, the cross-sectional study design precluded any inference of causality in the reported association between IS and DNAm levels. Third, there is a lack of replication in the meta-analysis. Fourth, we cannot draw conclusions about the effect of these methylation changes in gene expression because we lack gene expression data.

DNAm changes in 22 CpGs play a role in IS pathophysiology. They may orchestrate gene expression that contributes to IS and/or offer a biomarker of IS and vascular risk factors. Further investigation is required to assess the functional effect of DNAm changes and establish causality.

## ACKNOWLEDGEMENTS

We are grateful to all neurologists, nurses, and residents from the stroke unit and to technical staff from the IMIM who helped in performing this study. Elaine M. Lilly, PhD, provided English language assistance.

## Abbreviations

CE: Cardioembolic (stroke etiology)
DNAm: DNA methylation
EWAS: Epigenome-Wide Association Study
GWAS: Genome-Wide Association Study
IS: Ischemic Stroke
LAA: Large-Artery Atherosclerosis (stroke etiology)
MVPs: Methylation-Variable Positions
SNPs: Single Nucleotide Polymorphisms
SVD: Small Vessel Disease (stroke etiology)
TOAST: Trial of Org 10172 in Acute Stroke Treatment
UND: Undetermined (stroke etiology)

