## Supplemental Figures for "Identification of 20 novel loci associated to ischemic stroke. Epigenome-Wide Association Study"

**Supplementary Figure S1.** Region local plots showed the zoom-in view of the 22 MVPs replicated and other MVPs in the region was plotted with $-\log_{10}P$ values (left $y$-axis). Chr, chromosome ; CI, confidence interval; cM, centimorgan; and Mb, megabases.

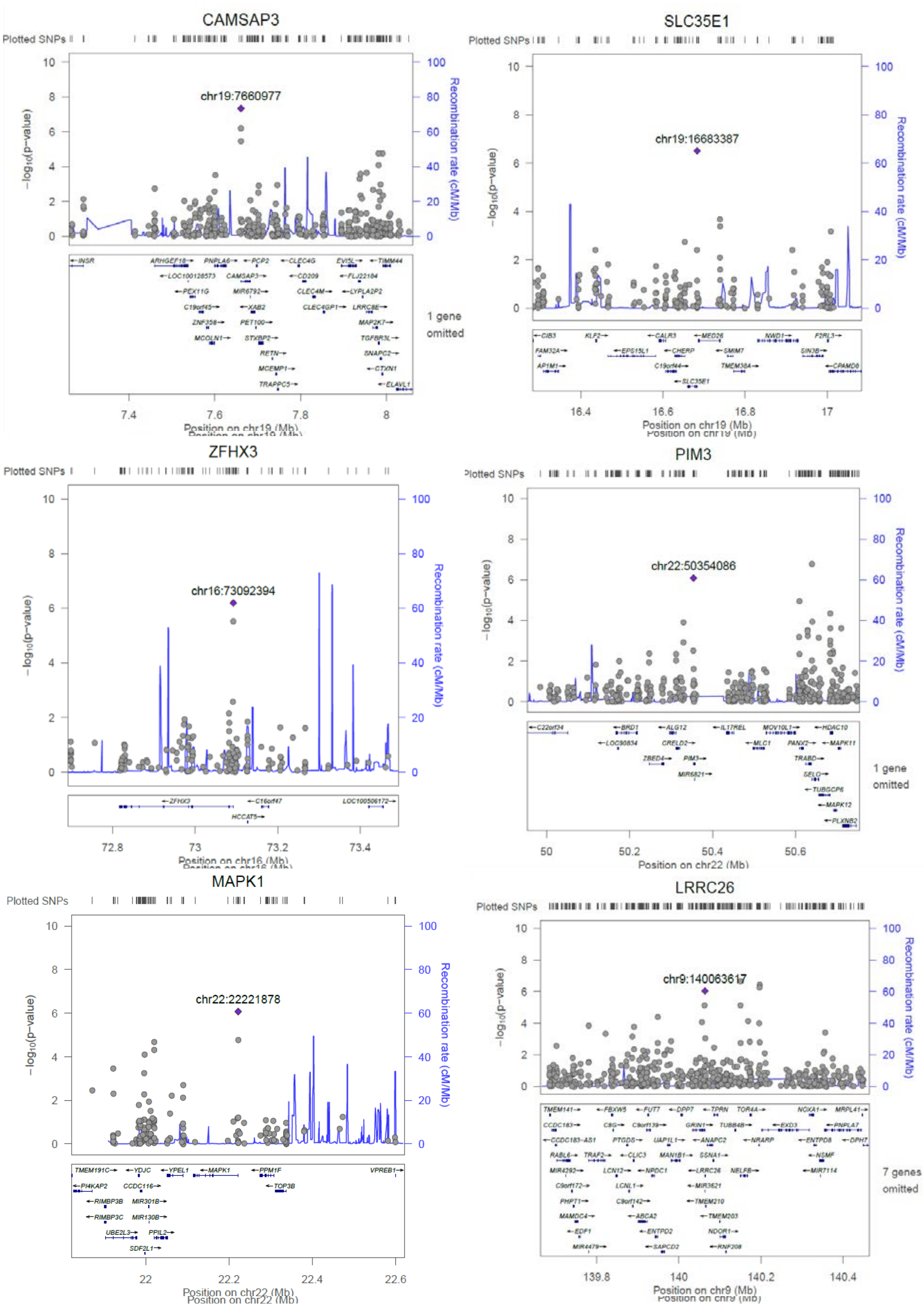

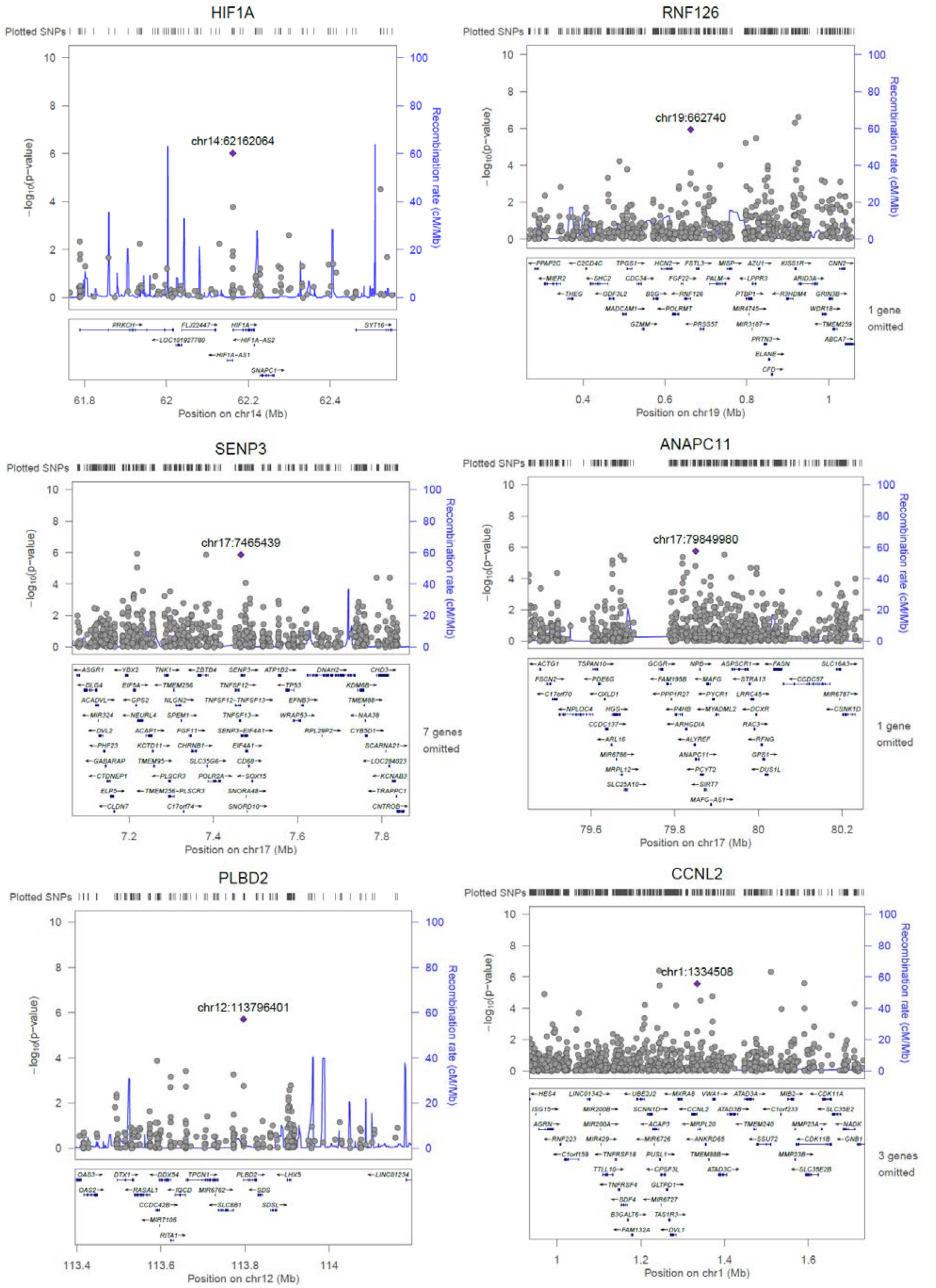
HIF1A
Plotted SNPs
chr14:62162064
−log10(p−value)
Recombination rate (cM/Mb)
Position on chr14 (Mb)
RNF126
chr19:662740
1 gene omitted
Position on chr19 (Mb)
SENP3
chr17:7465439
7 genes omitted
Position on chr17 (Mb)
ANAPC11
chr17:79849980
1 gene omitted
Position on chr17 (Mb)
PLBD2
chr12:113796401
Position on chr12 (Mb)
CCNL2
chr1:1334508
3 genes omitted
Position on chr1 (Mb)

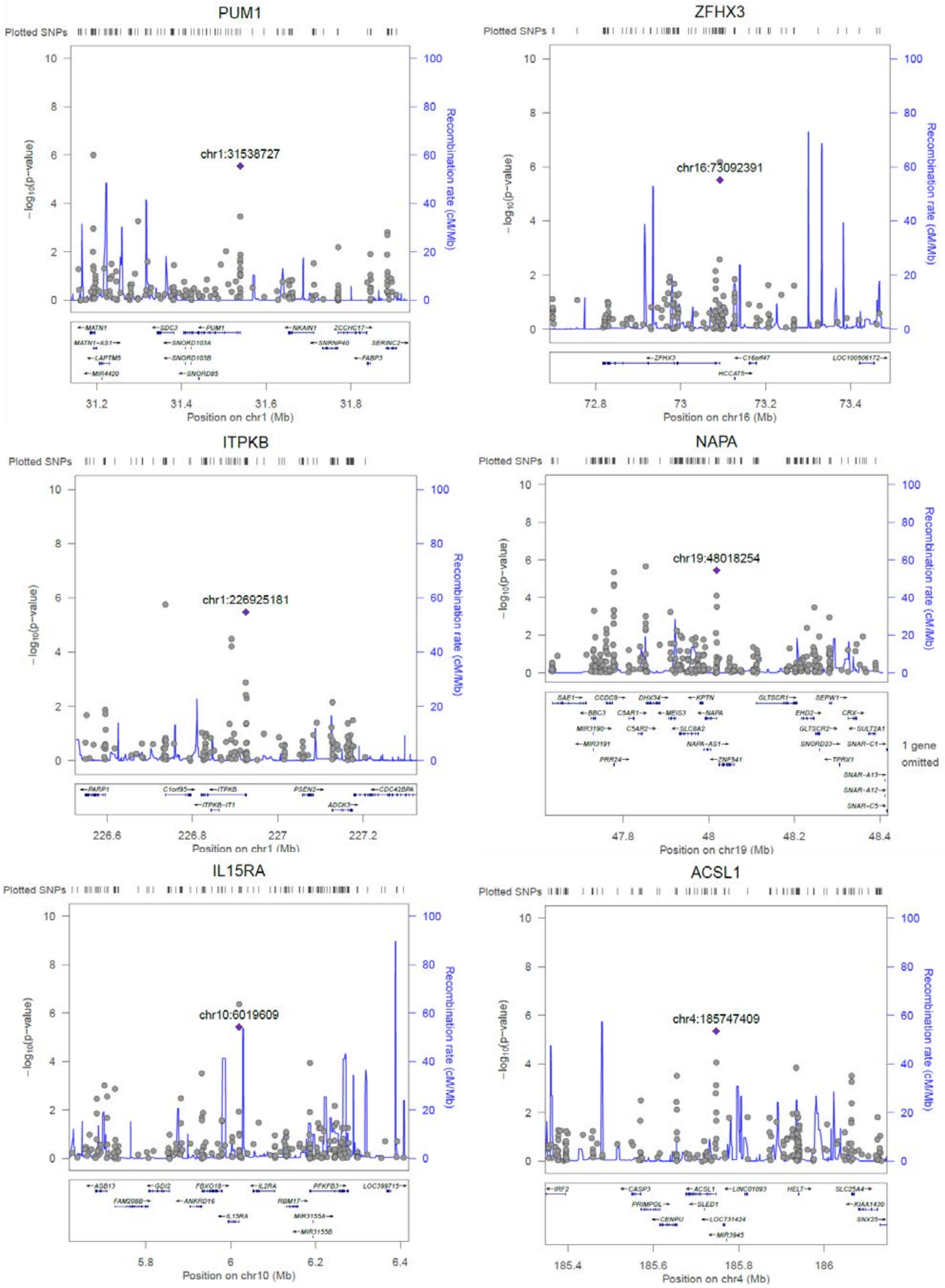
PUM1
Plotted SNPs
chr1:31538727
−log10(p−value)
Recombination rate (cM/Mb)
Position on chr1 (Mb)
ZFHX3
Plotted SNPs
chr16:73092391
−log10(p−value)
Recombination rate (cM/Mb)
Position on chr16 (Mb)
ITPKB
Plotted SNPs
chr1:226925181
−log10(p−value)
Recombination rate (cM/Mb)
Position on chr1 (Mb)
NAPA
Plotted SNPs
chr19:48018254
−log10(p−value)
Recombination rate (cM/Mb)
1 gene omitted
Position on chr19 (Mb)
IL15RA
Plotted SNPs
chr10:6019609
−log10(p−value)
Recombination rate (cM/Mb)
Position on chr10 (Mb)
ACSL1
Plotted SNPs
chr4:185747409
−log10(p−value)
Recombination rate (cM/Mb)
Position on chr4 (Mb)

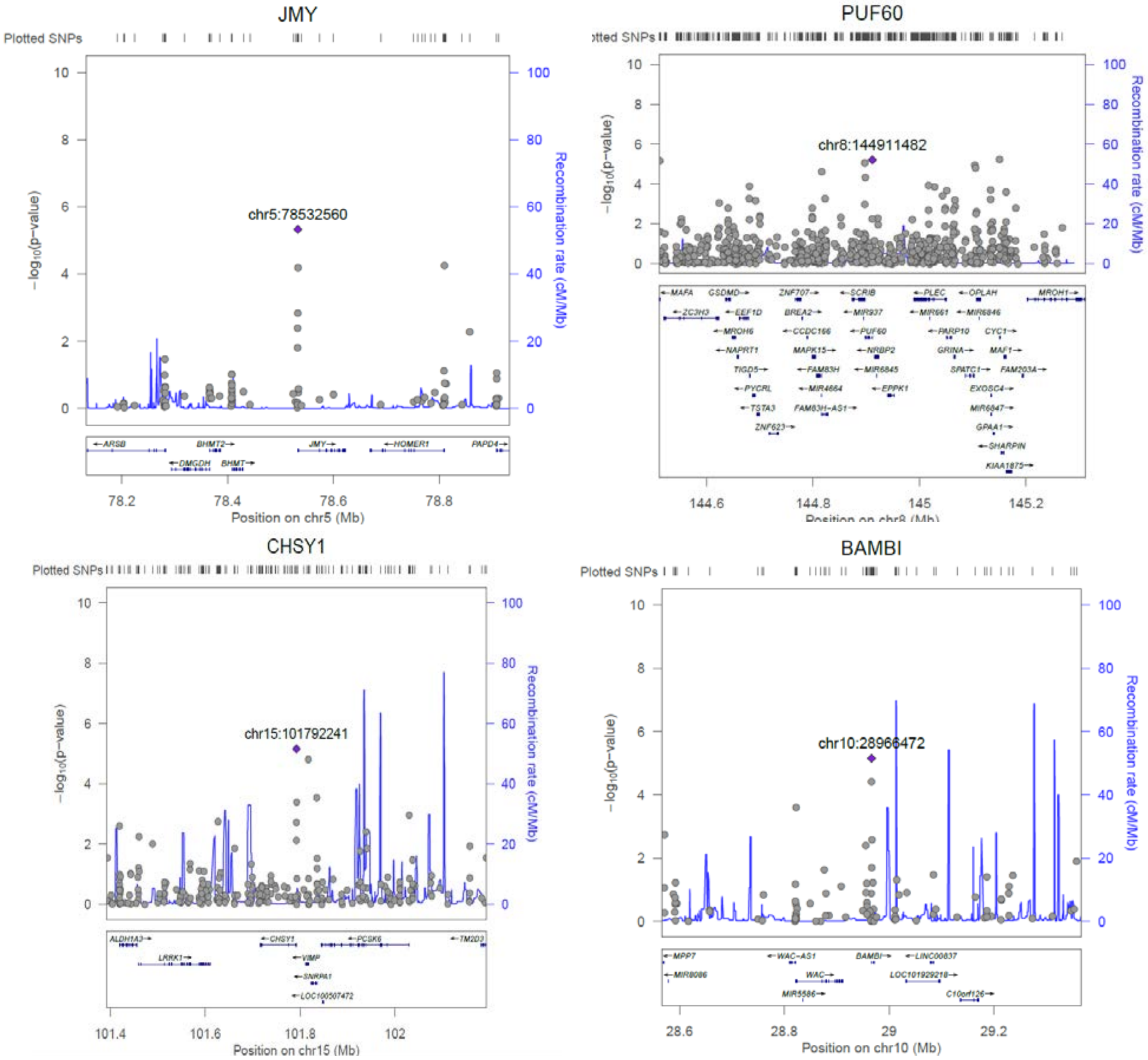
JMY
Plotted SNPs
chr5:78532560
−log10(p−value)
Recombination rate (cM/Mb)
Position on chr5 (Mb)
PUF60
Plotted SNPs
chr8:144911482
−log10(p−value)
Recombination rate (cM/Mb)
Position on chr8 (Mb)
CHSY1
Plotted SNPs
chr15:101792241
−log10(p−value)
Recombination rate (cM/Mb)
Position on chr15 (Mb)
BAMBI
Plotted SNPs
chr10:28966472
−log10(p−value)
Recombination rate (cM/Mb)
Position on chr10 (Mb)

**Supplementary Figure S2.** Genemania network. Interactions between the 21 loci enclosing validated CpGs and known loci associated to stroke.

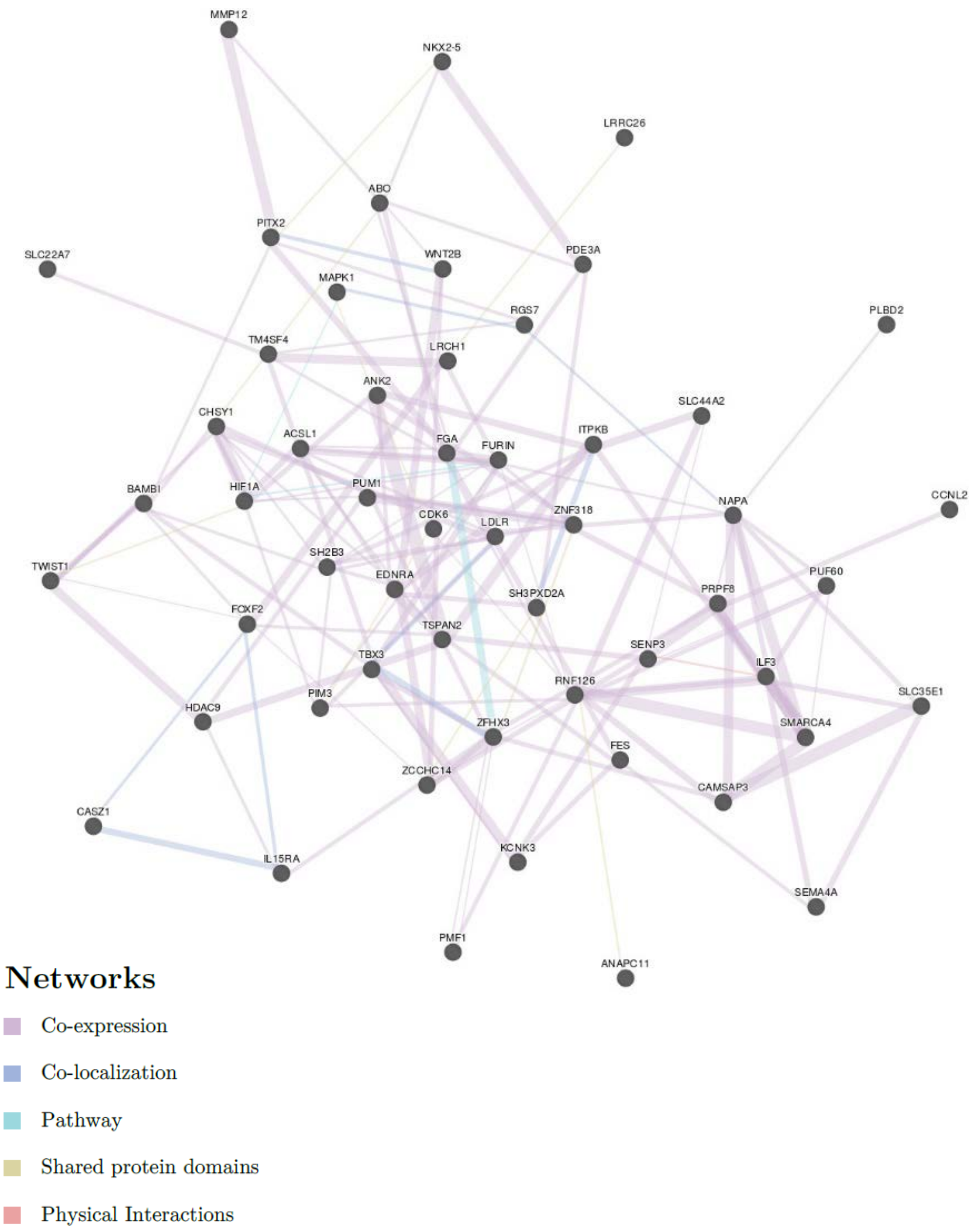

**Supplementary Figure S3.** GTEx tissue expression of the 21 loci enclosing validated. TPM, Transcripts Per Million.

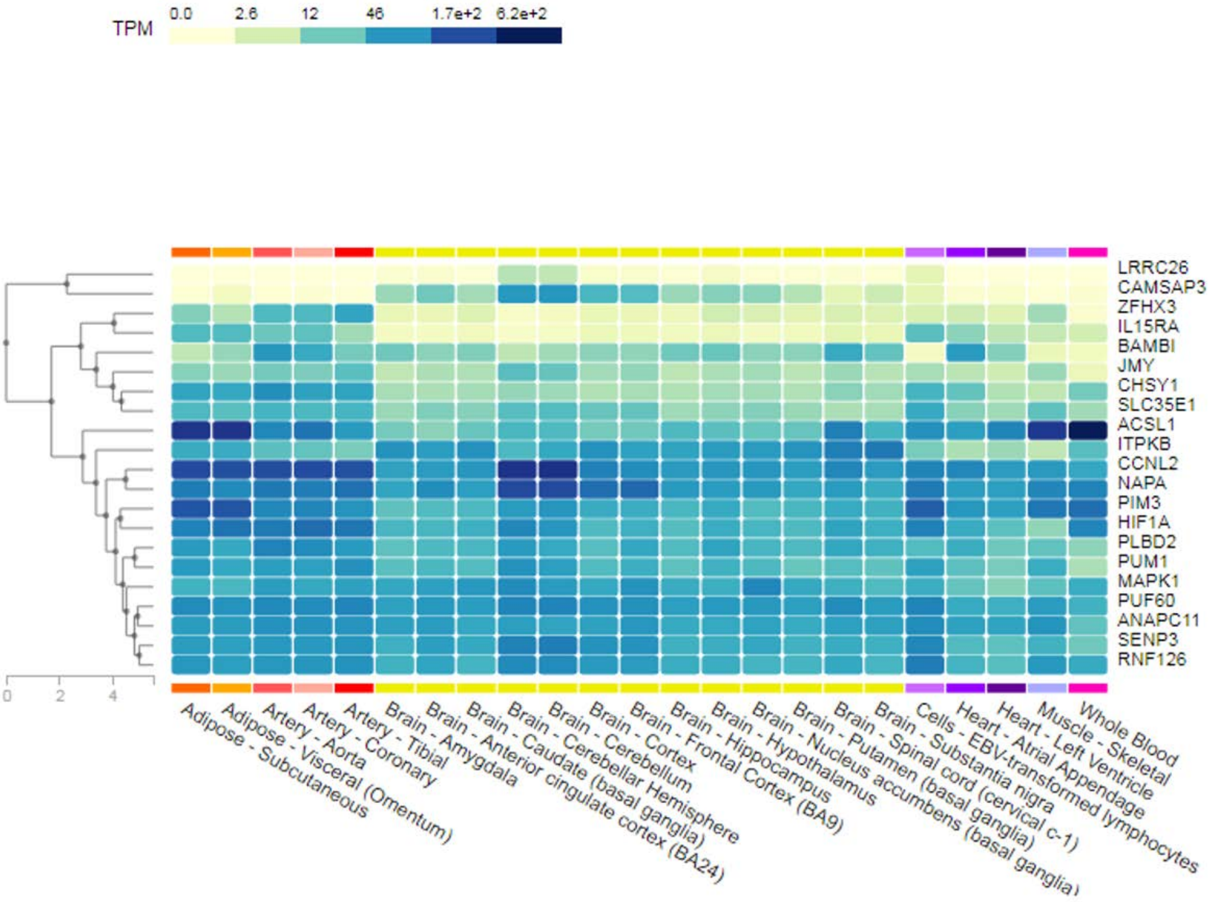

**Supplementary Figure S4.** Manhattan plots showing the distribution of the p-values of the associations between methylation probes in ischemic stroke patient analysis and QQ plot. (A) Large-Artery Atherosclerosis, (B) Cardiembolic ; (C) Small vessel diseases; (D) Undetermined subtypes.

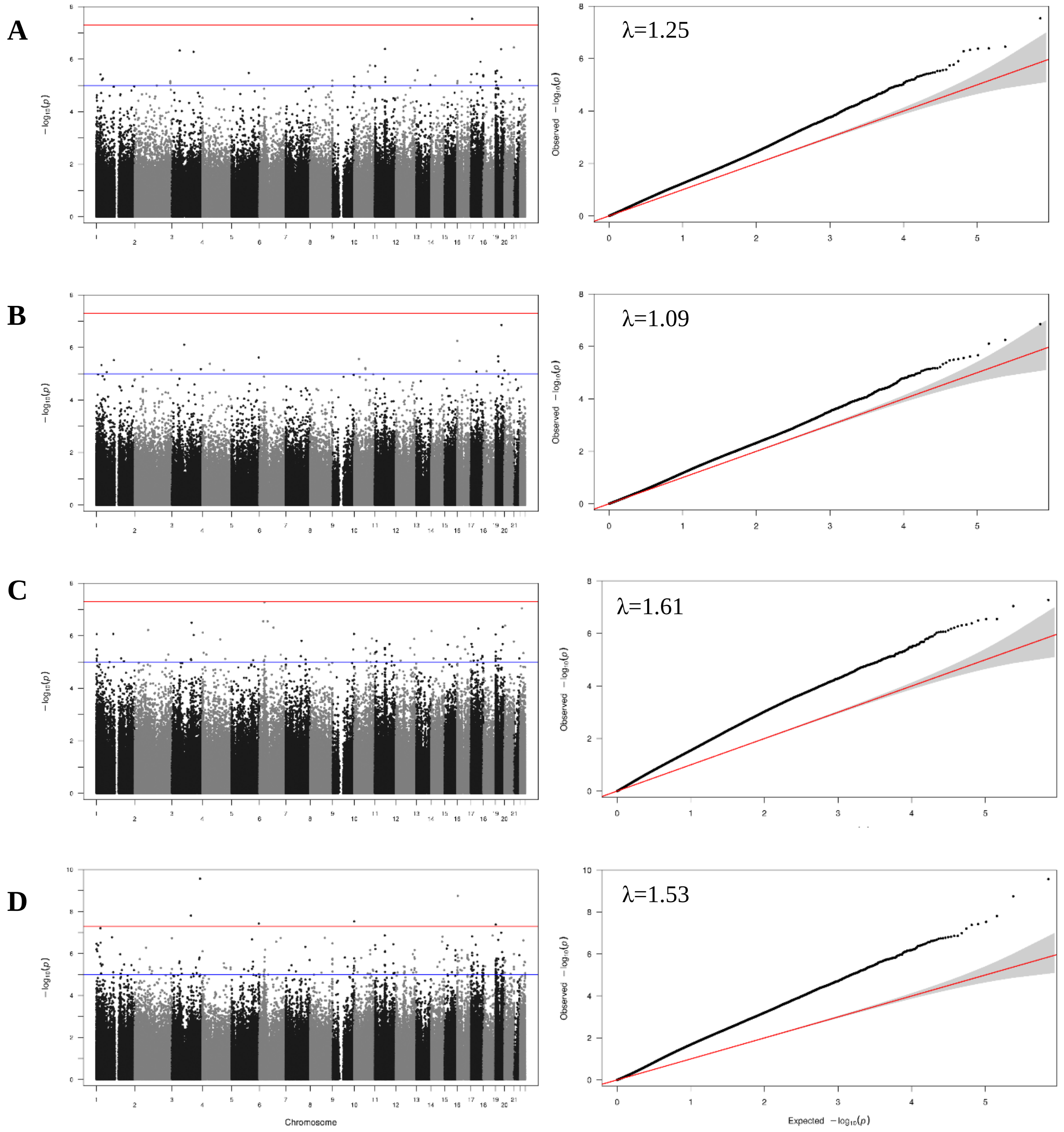
