## Supplemental Material for "Identification of 20 novel loci associated to ischemic stroke. Epigenome-Wide Association Study"

**Identification of 20 novel loci differentially methylation levels associated to ischemic stroke. Epigenome-Wide Association analysis**

Carolina Soriano-Tárraga[1] PhD, Eva Giralt-Steinhauer[1] MD, PhD, Uxue Lazcano[1] Msc, Carla Avellaneda-Gómez[1] MD, Ángel Ois[1] MD, PhD, Ana Rodríguez-Campello[1] MD, PhD, Elisa Cuadrado-Godia[1] MD, PhD, Alejandra Gomez-Gonzalez[1] MD, Alba Fernández-Sanlés[2] Msc, Roberto Elosua[2,3] MD, PhD, Israel Fernández-Cadenas[4] PhD, Natalia Cullell[4,5] Msc, Joan Montaner[6,7] MD, PhD, Sebastian Moran[8] PhD, Manel Esteller[9,10,11,12] MD, PhD, Jordi Jiménez-Conde[1] MD, PhD, and Jaume Roquer[1] MD, PhD

1. Neurovascular Research Group, Department of Neurology of Hospital del Mar-IMIM (Institut Hospital del Mar d'Investigacions Mèdiques); Universitat Autònoma de Barcelona/DCEXS-Universitat Pompeu Fabra, Barcelona, Spain.
2. Cardiovascular Epidemiology and Genetics Research Group, IMIM (Hospital del Mar Medical Research Institute), Barcelona, Spain.
3. CIBER de Enfermedades Cardiovasculares, Barcelona, Spain; Medicine Department, Medical School, University of Vic-Central University of Catalonia (UVic-UCC), Vic, Spain.
4. Stroke Pharmacogenomics and Genetics Group, Institut de Recerca Hospital de la Santa Creu i Sant Pau, Barcelona, Spain.
5. Neurology. Hospital Universitari MútuaTerrassa/Fundacio Docència i Recerca MútuaTerrassa, Spain
6. Neurovascular Research Laboratory, Vall d'Hebron Institute of Research (VHIR), Universitat Autònoma de Barcelona, Spain.
7. Institute de Biomedicine of Seville, IBiS/Hospital Universitario Virgen del Rocío/CSIC/University of Seville & Department of Neurology, Hospital Universitario Virgen Macarena, Seville, Spain.
8. Cancer Epigenetics and Biology Program (PEBC), Bellvitge Biomedical Research Institute (IDIBELL), L'Hospitalet, Barcelona, Spain.
9. Josep Carreras Leukaemia Research Institute (IJC), Badalona, Barcelona, Spain.
10. Centro de Investigacion Biomedica en Red Cancer (CIBERONC), Madrid, Spain.
11. Institucio Catalana de Recerca i Estudis Avançats (ICREA), Barcelona, Spain.
12. Physiological Sciences Department, School of Medicine and Health Sciences, University of Barcelona (UB), Barcelona, Spain.

**SUPPLEMENTAL MATERIAL**

1. Detailed Methods

2. Supplemental Figures and Figure Legends S1-4.

3. Supplemental Tables S1-23.

4. Supplemental References

### DETAILED METHODS

#### Discovery stage

The cases in the discovery sample (N=401, 183 controls/218 IS), MAR_1 sample, consisted of ischemic stroke (IS) patients were recruited in Hospital del Mar in Barcelona, Spain, from 2012 to 2015. It is a subset from those enrolled in BasicMar Register (Ministerio de Sanidad y Consumo, Instituto de Salud Carlos III; FIS No. PI051737), an ongoing prospective registry of stroke patients [1].

The BasicMar Register prospectively recruited all consenting patients who were admitted to our hospital from 2005 to 2019) with a diagnosis of stroke fulfilling World Health Organization criteria. Inclusion criteria in BASICMAR samples were as follows: (1) IS, (2) brain imaging with CT or MRI, (3) availability of the clinical data supporting the assigned stroke subtype according to TOAST classification[2], and (4) absence of intracranial hemorrhage, neoplasms, demyelinating and autoimmune diseases, and vasculitis. All patients were assessed and classified by a neurologist and were included in the study by consecutive order of recruitment.

Control samples (N=183) were obtained from Girona Heart Registry (REGICOR, which stands for REgistre GIroni del COR), a population-based cohort recruited in the province of Girona, in northeast Spain, about 100 km from Hospital del Mar (Barcelona) [3]. This register includes a randomized representative sample of men and women of the province of Girona. We used follow-up data from a population-based cohort originally enrolled in 2003-2005 (n=6352; response rate, 71.5%) from towns that represent the urban and rural diversity of Girona Province[3]. During 2009–2013, participants still residing in these towns were invited to participate in a follow-up visit; institutionalized residents were excluded. The response rate was 78.4%. A subsample of those attending their follow-up visit was selected as controls in this study. Inclusion criteria as controls: 1) No previous history of IS; 2) No previous history of acute myocardial infarction. All subjects were of European descent.

**Validation stage**

Two independent samples were used to replicate the results obtained in the discovery stage.

*Replication (1).* MAR_2 sample (N=226), it is a second subsample of IS patients (N=185) from those enrolled in the BasicMar Register, was used as one of the replication samples, patients recruited from 2009 to 2012. A total of 41 REGICOR samples were included as controls.

*Replication (2).* HVH sample (N=166), with 145 IS patients and 21 controls, was used as the second replication sample, recruited from 2009 to 2012 in Hospital Vall d'Hebron in Barcelona (Spain).

**Stroke Subtype Classification**

Using TOAST criteria, patients were classified into 4 categories: (1) large-artery atherosclerosis (LAA), (2) small-vessel disease (SVD), (3) cardioembolic (CE), and (4) stroke of undetermined etiology (UND). Diagnoses were based on clinical features and on data collected by methods such as brain imaging (CT/MRI) and cardiac imaging [2].

**Demographic and Vascular Risk Factor Variables**

In accordance with international guidelines, data on vascular risk factors analyzed were obtained from a direct interview of the patient, relatives and caregivers, and from medical records. Examinations were performed and standardized questionnaires administered during the hospitalization by a team of neurologists and reviewed by an additional neurologist.

We recorded age, sex, and vascular risk factors using a structured questionnaire, as follows: arterial hypertension (HT), defined as systolic blood pressure ≥140 mmHg or diastolic ≥90 mmHg recorded from more than 2 measurements previous to the acute event, a physician's diagnosis, or use of medication; hyperlipidemia (HL), defined as a physician's diagnosis, use of medication, serum cholesterol concentration >220 mg/dL, low-density lipoprotein cholesterol >130 mg/dL, or serum triglyceride concentration >150 mg/dL; diabetes mellitus (DM), defined as evidence of two or more fasting blood glucose values ≥126 mg/dl, use of diabetes medication, or a physician's diagnosis; coronary heart disease (CHD), defined as documented history of angina pectoris or myocardial infarction; atrial fibrillation (AF) (documented history or diagnosis during hospitalization); and self-reported smoking habit. During hospitalization, body mass index (BMI), initial stroke severity (measured by the National Institutes of Health Stroke Scale (NIHSS)), smoking status were recorded and TOAST criteria were used to classify IS subtype [2], according to standardized protocol.

**Peripheral Blood Collection and DNA Extraction**

IS DNA samples were extracted from whole peripheral blood collected in 10 mL EDTA tubes at hospital arrival, in the acute phase of the stroke (maximum within 12 hours of symptoms onset). The Chemagic Magnetic Separation Module I system (Chemagen), The Autopure LS (Qiagen) and Gentra Puregene Blood Kit (Qiagen, Hilden, Germany) were used for DNA isolation in BASICMAR samples. The Autopure LS (Qiagen) was used for DNA isolation in the REGICOR sample. The Gentra Puregene Blood Kit (Qiagen, Hilden, Germany) was used in the HVH sample.

DNA extractions stored together at −20°C. DNA concentrations were quantified using Picogreen assay and Nanodrop technology. The quality of DNA samples was visualized in agarose gels.

**Array-based DNA methylation analysis**

Genomic DNA (1 µg) was bisulfite converted using EZ-96 DNA Methylation Kit (Zymo Research, Orange, CA,USA) according to the manufacturer's procedure, with the alternative incubation conditions recommended when using the Illumina Methylation Assay.

Genome-wide DNA methylation of the discovery sample, MAR_1 and REGICOR samples, was assessed in the same Infinium MethylationEPIC Beadchip arrays (Illumina Netherlands, Eindhoven, Netherlands) following the manufacturer's protocol with no modifications. This array covers ~850.000 methylation CpG sites. The arrays were scanned with the Illumina HiScan SQ scanner. MAR_2 sample was assessed using the Illumina HumanMethylation450 Beadchip (Illumina Netherlands, Eindhoven, The Netherlands) following the manufacturer's protocol with no modifications. This array covers 485,577 methylation CpG sites in 99% of RefSeq genes (21,231 genes). The arrays were scanned with the Illumina HiScan SQ scanner. HVH sample was assessed using the Illumina HumanMethylation450 and Infinium MethylationEPIC Beadchip arrays (Illumina Netherlands, Eindhoven, The Netherlands)

**Data Pre-processing and Normalization**

Sample and CpG quality controls and the statistical analysis were performed as described in Soriano-Tarraga et al [4]. Only probes common in HumanMethylation450 Beadchip were analized.

*Sample quality control.* We used all the samples that had a detection rate over 95%. Then, we tested whether we could group samples by sex according to their DNA methylation levels on the X-chromosome using the *methylumi* R package [5]. Samples that were poorly performing in theses quality controls were excluded from further analysis.

*CpG quality control.* We excluded all probes that were represented by a bead count under 3 in at least 5% of the samples. CpG sites having 1 % of samples with a detection p-value greater than 0.05 were removed and cross-reactive probes were excluded [6] using the watermelon R package [7]. To avoid SNP (single-nucleotide polymorphism) effects on methylation measures and sex bias, we excluded CpG probes in close proximity to common SNPs and all probes associated to allosomal position.

Before analysis, methylation values were corrected for background values and then normalized by *Noob* using *minfi* Bioconductor package [8,9]. A total of 401 samples and 358,709 autosomal CpGs were analyzed. (**Supplementary Material, Table S1-2**)

We used commonly used algorithms to infer both white blood cell counts[10] and omics-related confounding[11] from DNA methylation data, which were subsequently included as covariates in the association analyses. We used the array annotations provided by Illumina to assign probes to the corresponding genes.

**Statistical Analysis**

Baseline characteristics were compared between IS and controls using t-test for continuous, and chi-squared for categorical variables. Continuous variables are presented as means and standard deviation (SD) or medians and interquartile ranges (IQR), and categorical variables as absolute values and percentages. For the bivariate analyses, baseline characteristics of the IS and controls were compared using *Student t-test* for continuous variables and $\chi 2$ test for categorical variables.

First, we analyzed the association between DNA methylation at all the individual CpG sites comparing controls vs IS samples. We included all the individuals and CpG sites that passed quality controls (**Supplementary Material, Table S1**). We analyzed for differences in methylation at the CpG sites or methylation-variable positions (MVPs), between two groups, controls and IS, using a multivariate lineal regression model adjusting by sex, age, array, slide, smoking status, NIHSS, HT, HL, DM, CHD, AF and cell count. The reason of adjusting by stroke severity (NIHSS) was to reduce as much as the possible effects of collecting the blood samples during the acute phase of IS.

The analyses were performed using the R statistical package, version 3.5.2 [12]. The following packages were used: *minfi* and *limma* [9,13]. The global significance level of 0.05 was corrected for multiple comparisons, a p-value<$1.39\times10^{-7}$ (0.05/358,709 CpgG sites), was established to define the statistically significant differences and a p-value<$1.39\times10^{-5}$ as nominally significant in the discovery stage. The top 500 CpGs in were used in the replication stage, arbitrary p-value $\leq7.1\times10^{-6}$. We also used LocusZoom (http://csg.sph.umich.edu/locuszoom) to generate regional association plots [14].

**Replication stage and Meta-analysis**

Top 500 CpGs in the discovery stage and that were represented in both arrays (450k and EPIC) were included for replication and meta-analysis (N=793) into weighted z-score meta-analyses using METAL [15]. The global significance level of 0.05 was corrected for multiple comparisons, in MAR_2 sample p-value$\leq1.0\times10^{-4}$ (0.05/500 CpG sites) and HVH sample p-value$\leq1.09\times10^{-4}$ (0.05/451 CpG sites) were used to define statistically significant differences. In the meta-analysis p-value<$1.39\times10^{-7}$ was established to define the statistically significant differences.

**Pathway analysis**

We used the GeneMANIA (http://pages.genemania.org/) algorithm to look for relationships among candidate genes and known IS-related genes [16] by searching within multiple publicly available biological datasets. These datasets include protein-protein, protein-DNA and genetic interactions, pathways, reactions, gene and protein expression data, protein domains, and phenotypic screening profiles.

The significant loci were uplodaded to FUMA (v1.3.5) to annotate GWAS significant variants from GWAS catalog (e96_r2019-05-03), gene expression (GTEx/v6/gtex_v6_ts_avg_log2RPKM, GTEx/v6/gtex_v6_ts_general_avg), Ensemble (v92) and pathways MsigDB (v6.2). Gene-based analysis was also performed by FUMA [17]. Moreover, PheGenI (https://www.ncbi.nlm.nih.gov/gap/phegeni) and DAVID 6.8 (https://david.ncifcrf.gov/tools.jsp) were used to find GWAS associations, and to asses enrichment analysis, respectivelly [18]

**Standard Protocol Approvals, Registrations, and Patients Consents**

Local ethics committees, CEIC-Parc de Salut Mar and the Ethics Committee of the Vall d'Hebron Hospital, Barcelona, approved the study. Written Informed consent was provided by all participants or their approved proxy. The principles expressed that the Declaration of Helsinki and relevant national legislation were followed.

**Data Availability Policy**

All data generated from this study is included in the main manuscript and its supplementary information file.

### SUPPLEMENTAL FIGURES AND FIGURE LEGENDS

**Supplemental Figure S1.** Region local plots showed the zoom-in view of the 22 MVPs replicated and other MVPs in the region was plotted with $-\log_{10}P$ values (left *y*-axis). Chr, chromosome; CI, confidence interval; cM, centimorgan; and Mb, megabases.

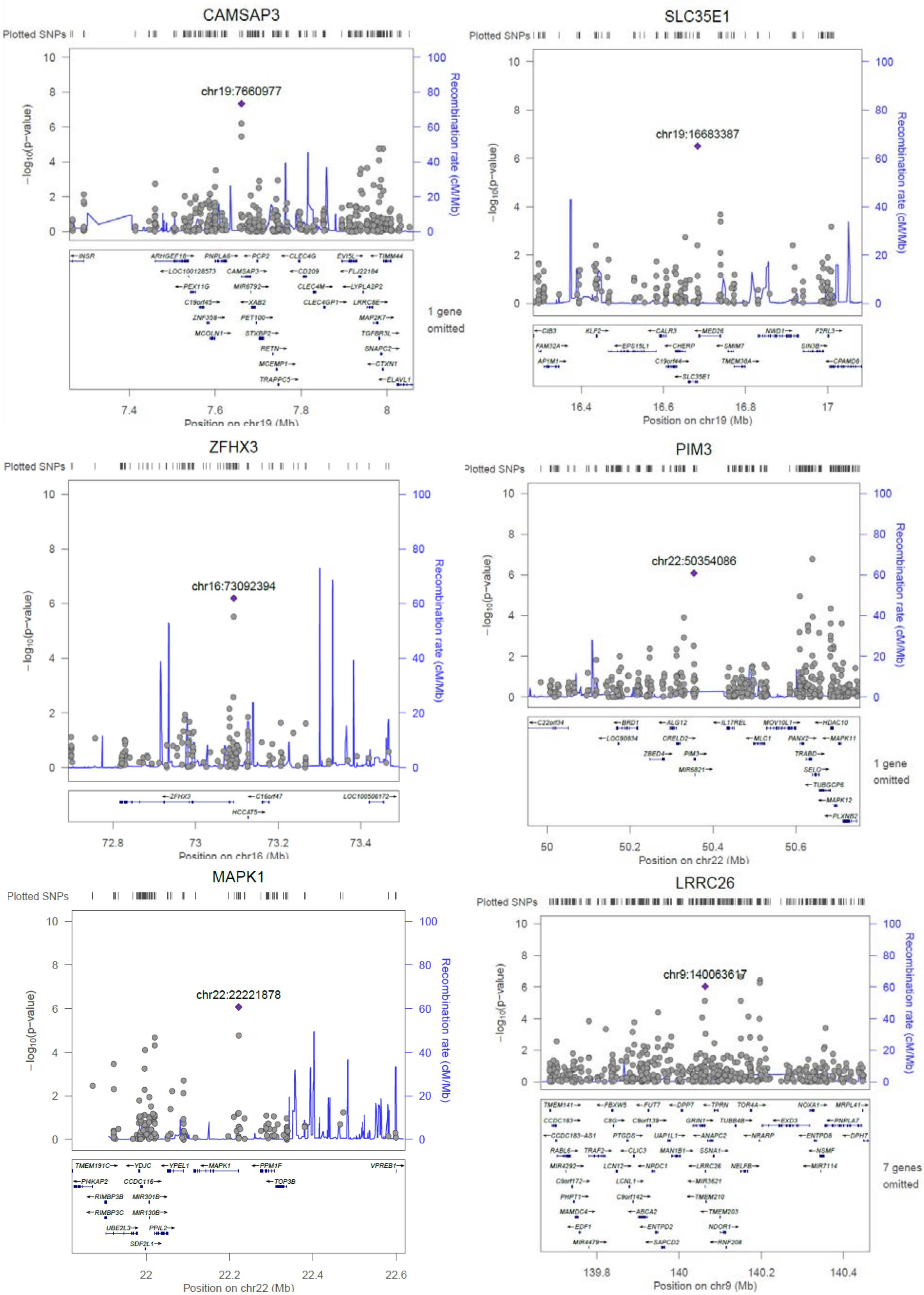

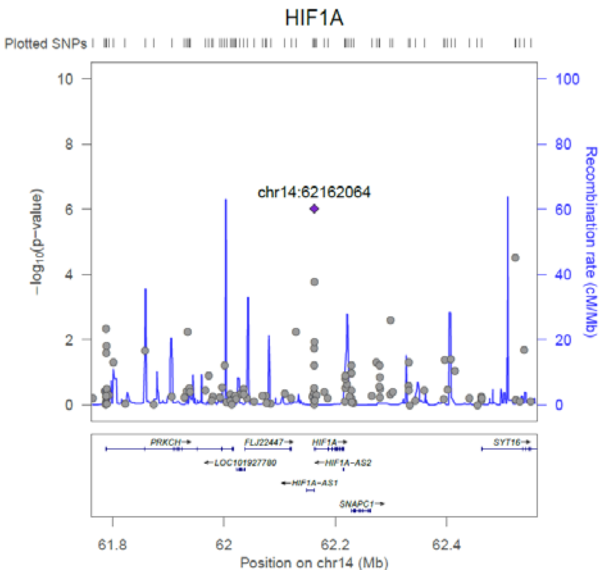
HIF1A
Plotted SNPs
chr14:62162064
−log10(p−value)
Recombination rate (cM/Mb)
Position on chr14 (Mb)

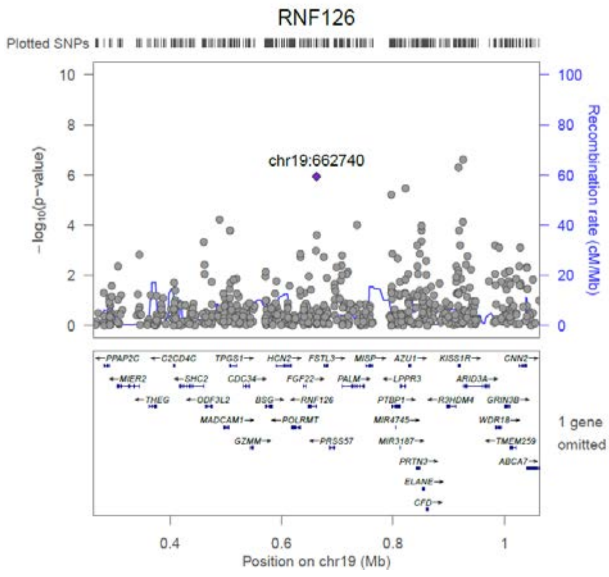
RNF126
Plotted SNPs
chr19:662740
−log10(p−value)
Recombination rate (cM/Mb)
1 gene omitted
Position on chr19 (Mb)

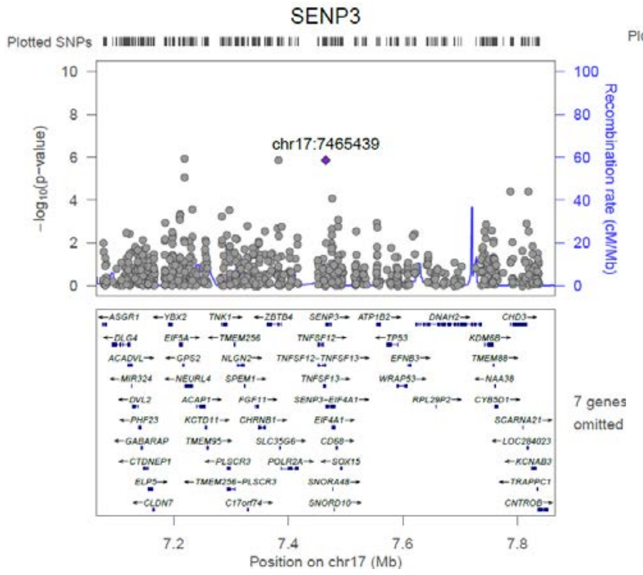
SENP3
Plotted SNPs
chr17:7465439
−log10(p−value)
Recombination rate (cM/Mb)
7 genes omitted
Position on chr17 (Mb)

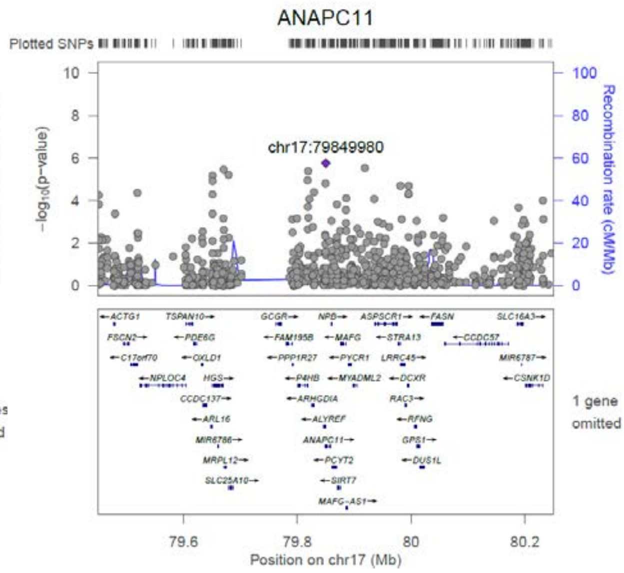
ANAPC11
Plotted SNPs
chr17:79849980
−log10(p−value)
Recombination rate (cM/Mb)
1 gene omitted
Position on chr17 (Mb)

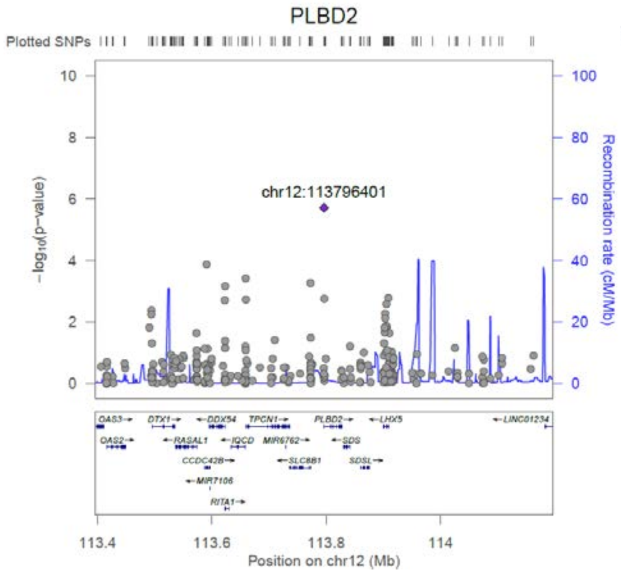
PLBD2
Plotted SNPs
chr12:113796401
−log10(p−value)
Recombination rate (cM/Mb)
Position on chr12 (Mb)

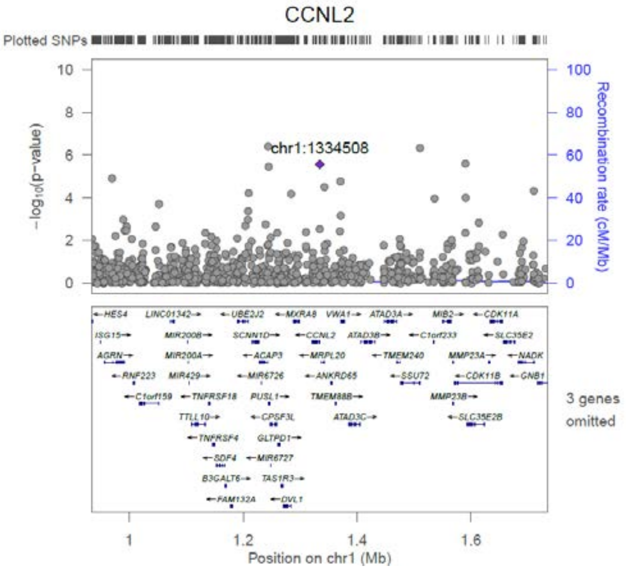
CCNL2
Plotted SNPs
chr1:1334508
−log10(p−value)
Recombination rate (cM/Mb)
3 genes omitted
Position on chr1 (Mb)

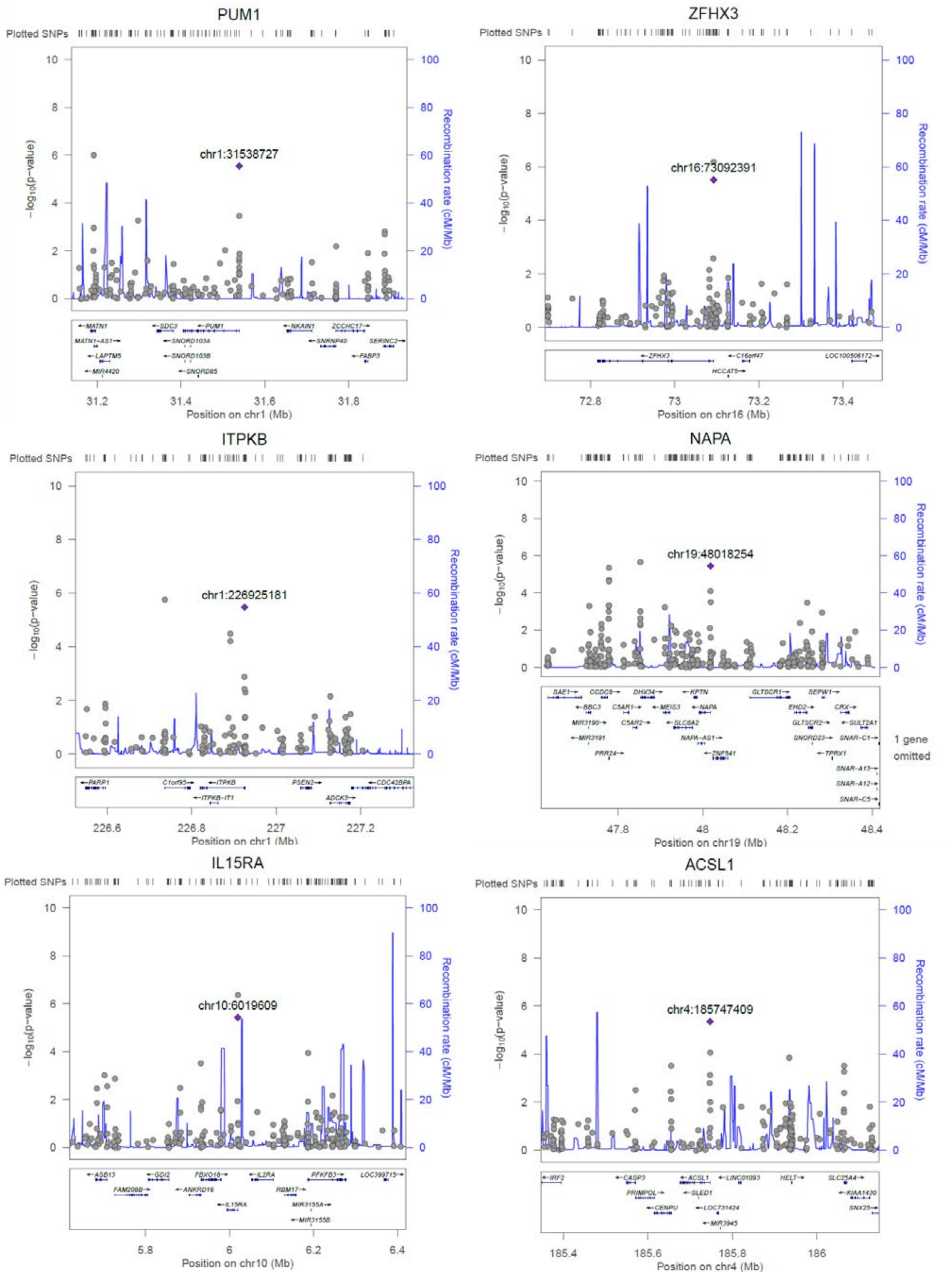
PUM1
Plotted SNPs
chr1:31538727
-log10(p-value)
Recombination rate (cM/Mb)
Position on chr1 (Mb)
ZFHX3
Plotted SNPs
chr16:73092391
-log10(p-value)
Recombination rate (cM/Mb)
Position on chr16 (Mb)
ITPKB
Plotted SNPs
chr1:226925181
-log10(p-value)
Recombination rate (cM/Mb)
Position on chr1 (Mb)
NAPA
Plotted SNPs
chr19:48018254
-log10(p-value)
Recombination rate (cM/Mb)
1 gene omitted
Position on chr19 (Mb)
IL15RA
Plotted SNPs
chr10:6019609
-log10(p-value)
Recombination rate (cM/Mb)
Position on chr10 (Mb)
ACSL1
Plotted SNPs
chr4:185747409
-log10(p-value)
Recombination rate (cM/Mb)
Position on chr4 (Mb)

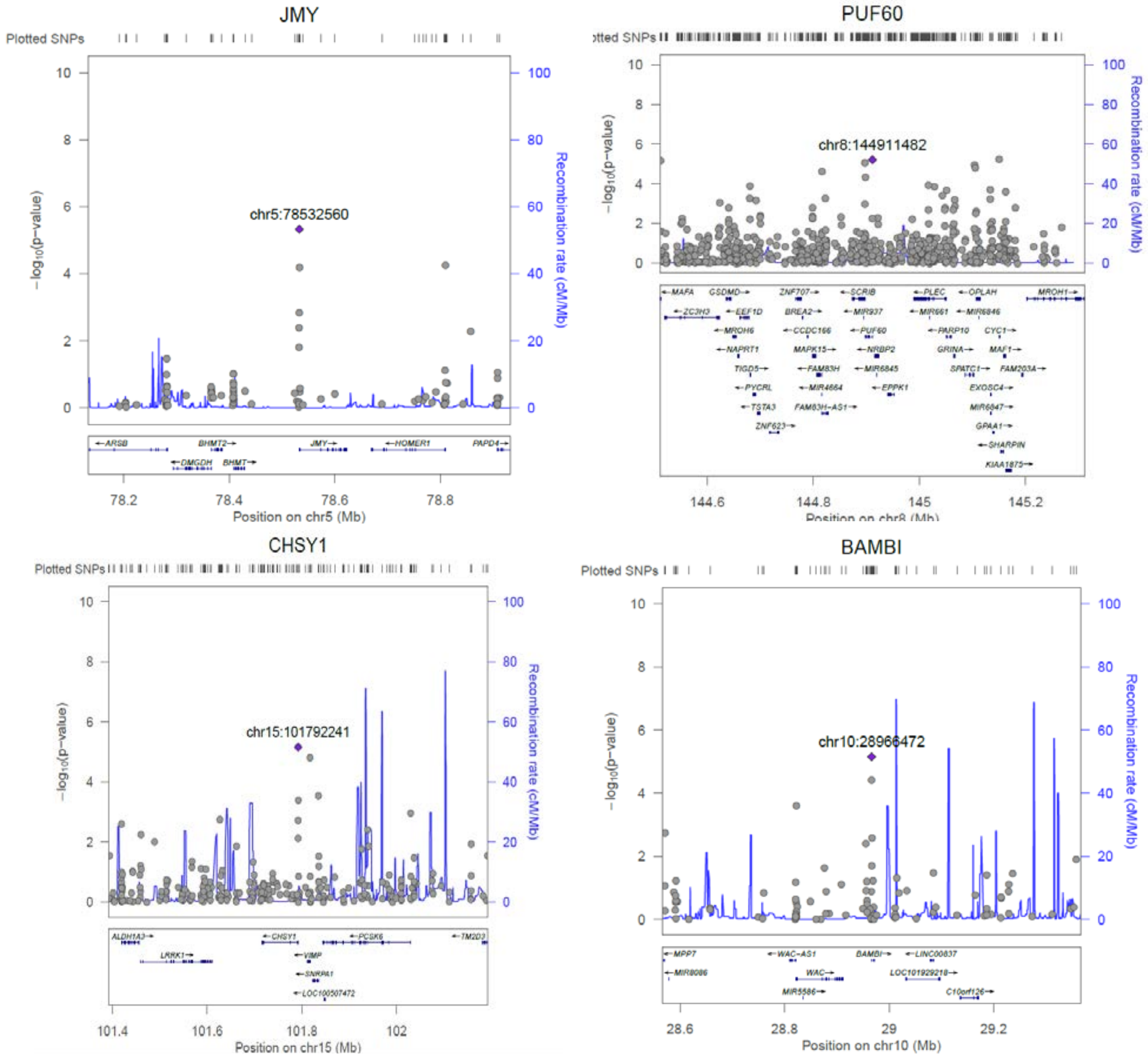
JMY
Plotted SNPs
chr5:78532560
−log10(p−value)
Recombination rate (cM/Mb)
Position on chr5 (Mb)
PUF60
chr8:144911482
Position on chr8 (Mb)
CHSY1
Plotted SNPs
chr15:101792241
Position on chr15 (Mb)
BAMBI
Plotted SNPs
chr10:28966472
Position on chr10 (Mb)

**Supplemental Figure S2.** Genemania network. Interactions between the 21 loci enclosing validated CpGs and known loci associated to stroke.

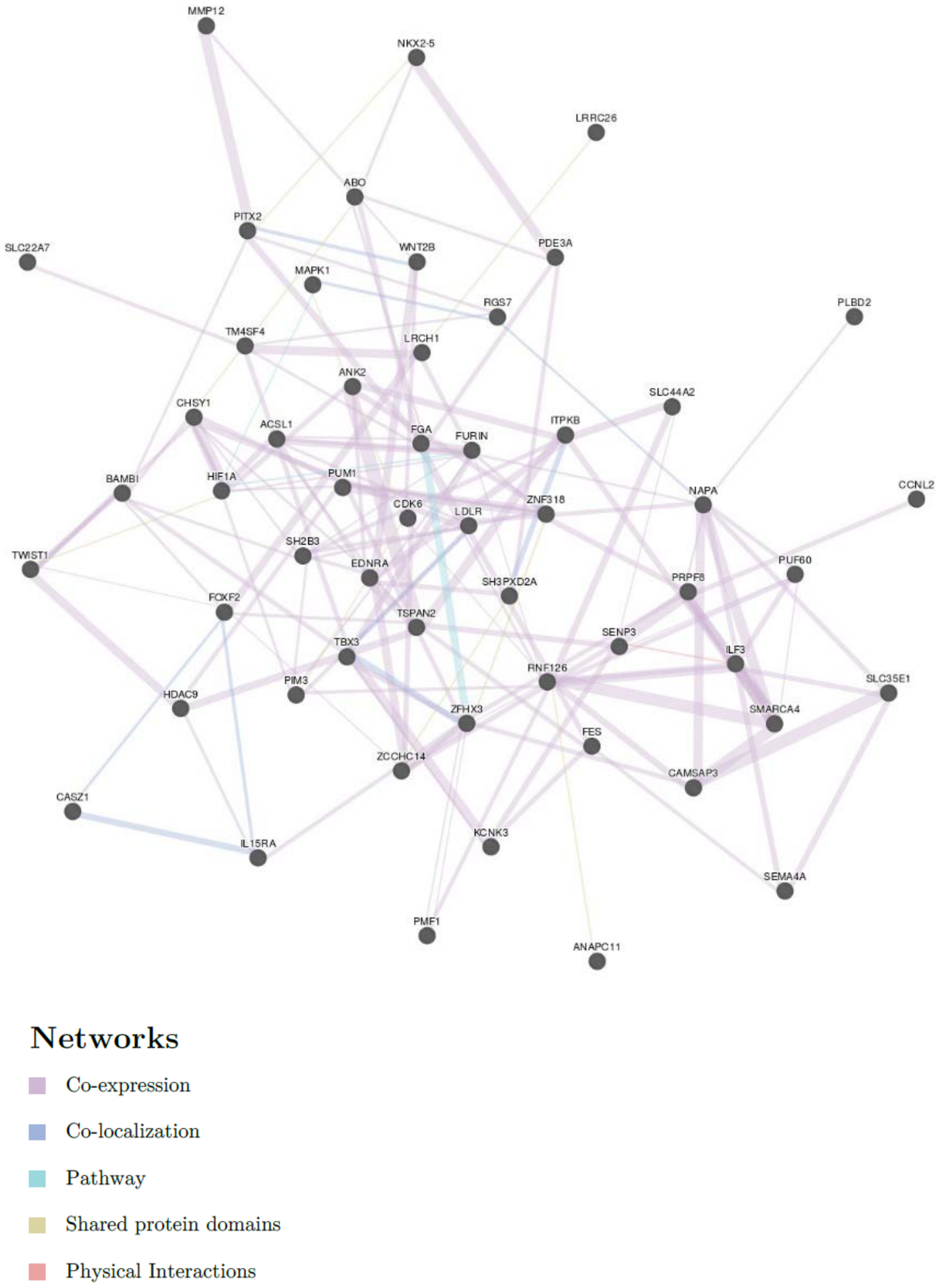

**Supplemental Figure S3.** GTEx tissue expression of the 21 loci enclosing validated. TPM, Transcripts Per Million.

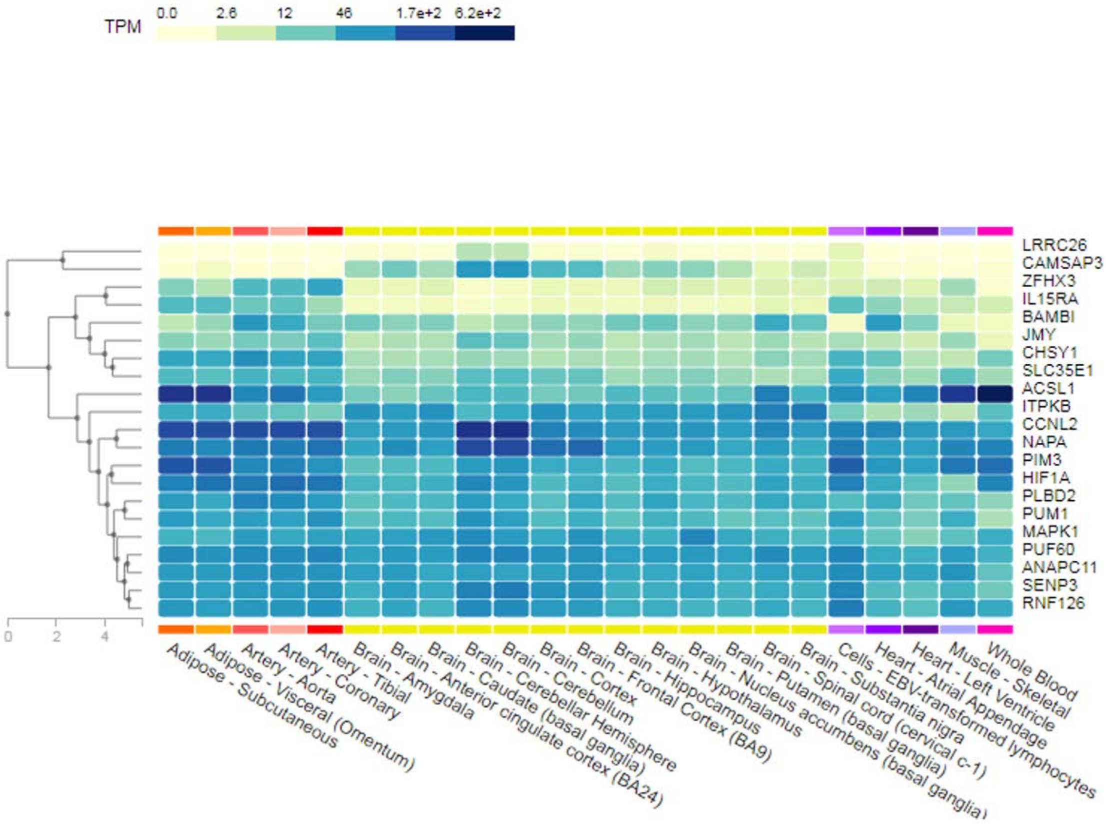

**Supplemental Figure S4**. Manhattan plots showing the distribution of the p-values of the associations between methylation probes in ischemic stroke patient analysis and QQ plot. (A)  Large-Artery Atherosclerosis, (B) Cardiembolic ; (C) Small vessel diseases; (D) Undetermined subtypes.

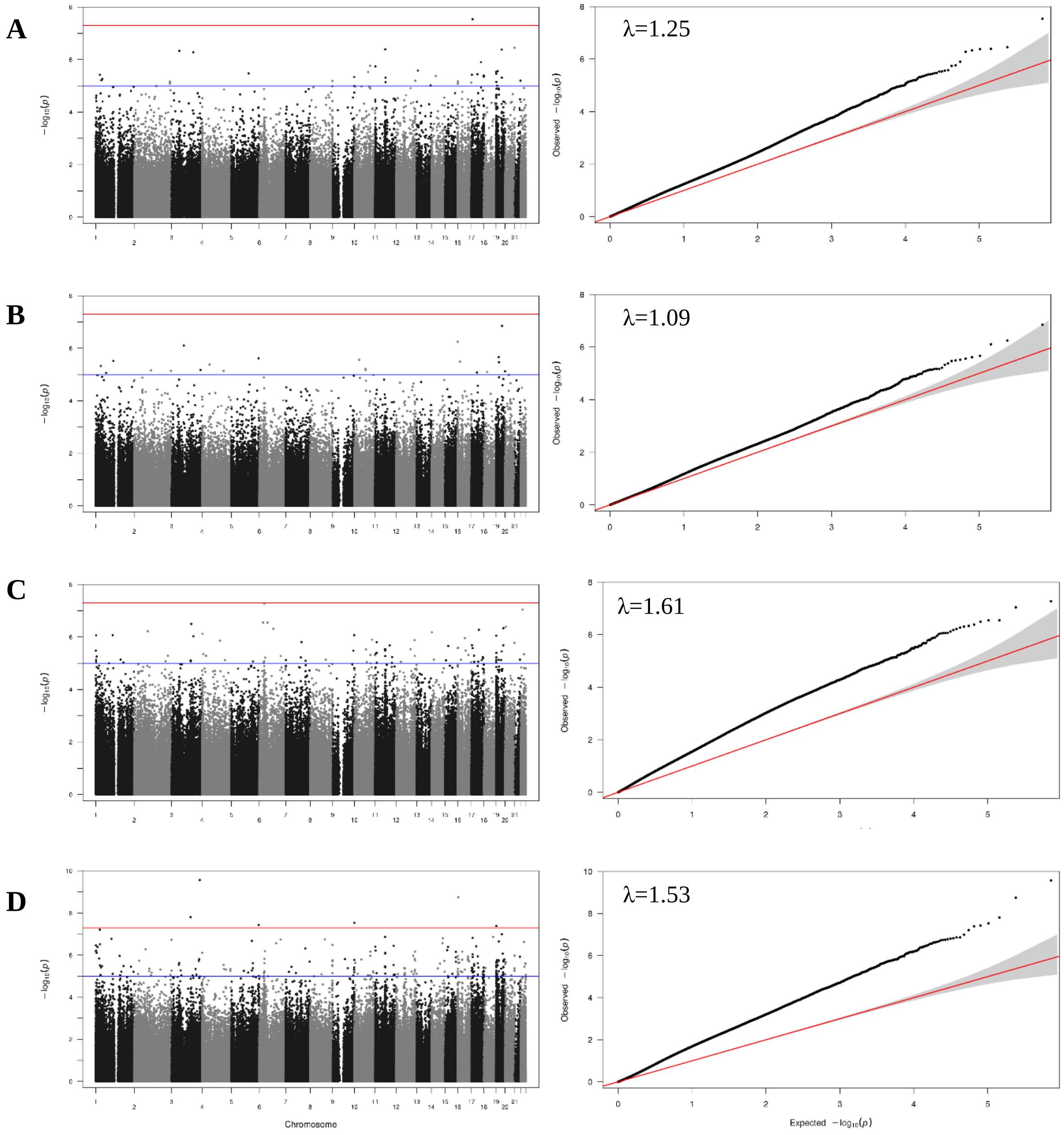

**SUPPLEMENTAL TABLES**

**Supplemental Table S1.** Infinium Methylation EPIC Beadchip quality control and selection of 450k probes**.** We assessed the quality control of the process, focusing separately on sample quality control and CpG site (i.e., probe) quality control, following a standardized pipeline. Table shows excluded samples and probes at each step. SNP, probe deleted containing genetic variant categories with minor allele frequency > 5 % at target CpG sites (CpG), single base extension sites of Type I probes (SBE) and overlapping the probe body (Probe).

| | **Discovery (MAR_1 + REGICOR)** |
|---|---|
| **Inicial numbers** | 835 samples<br>866,836 probes |
| **Removed Samples by QC** | 79 |
| detection rate over 95% | 3 |
| sex according X-chromosome | 76 |
| **Removed Probes by QC** | 160,720 |
| beadcount <3 in 5 % of samples and | 560 |
| sites having 1 % of samples with p-value > 0.05 | 13,765 |
| SNP | 98,803 |
| cross-reactive-probes | 28,030 |
| probes from Chromosomes X and Y | 19,627 |
| **Removed Samples by exclusion criteria** | 355 |
| **Removed Probes no common to 450K Beadchip** | 347,342 |
| **Final number of samples (%)** | 401 (48.0%) |
| **Final number of probes (%)** | 358,709 (41.4%) |

**Supplemental Table S2**. DNA methylation arrays used by samples and number of CpGs analyzed.

| **Samples** | **Illumina Platform** | **CpGs** | **N (controls/IS)** |
|---|---|---|---|
| **Discovery** | Methylation EPIC Beadchip.<br>Selection of 450k probes | 358,709 | 401 (183/218) |
| **Replication (1)** | Human Methylation450 Beadchip | 500 | 226 (41/185) |
| **Replication (2)** | Human Methylation450 Beadchip<br>Methylation EPIC Beadchip | 451 | 166 (21/145) |
| **Joint** | | | 793(245/548) |

**Supplemental Table S3. Discovery stage. Ischemic stroke patients vs controls.** CpG sites differentially methylated in relation to ischemic stroke. CpG id; Chr, chromosome location; genomic position; associated gene and observed mean β-values (standard deviation).

See Supplemental Tables, Excel file

**Supplemental Table S4.** Descriptive characteristics of the 21 genes where the 22 MVPs are.

| Gene | Traits and Functions | Reference |
|---|---|---|
| **CAMSAP3** | Calmodulin regulated spectrin associated protein family member 3. Maintain neuronal polarity. | 19 |
| **SLC35E1** | Solute carrier family 35 member E. Gene highly conserved | - |
| **ZFHX3** | Zinc finger homeobox 3. Associated to atrial fibrillation, ischemic stroke, Body mass Index and LDL cholesterol levels. | 16,20,21 |
| **PIM3** | Pim-3 proto-oncogene, serine/threonine kinaseProto-oncogene with serine/threonine kinase activity that can prevent apoptosis, promote cell survival and protein translation. Additionally to its role on tumorigenesis, can also negatively regulate insulin secretion, control of energy metabolism and cell growth. | 22 |
| **MAPK1** | Mitogen-activated protein kinase 1.Serine/threonine kinase which acts as an essential component of the MAP kinase signal transduction pathway. Transmission of signals in response to different stimuli such as ischemia or inflammation | 23–25 |
| **LRRC26** | Leucine rich repeat containing 26.Elevates channel voltage- and apparent Ca(2+) sensitivity in arterial myocytes to induce vasodilation. | 26 |
| **HIF1A** | Hypoxia inducible factor 1 subunit alpha. Master regulator of cellular and systemic homeostatic response to hypoxia by activating transcription of many genes, including those involved in energy metabolism, angiogenesis, apoptosis, among others to facilitate metabolic adaptation to hypoxia. | 27 |
| **RNF126** | Ring finger protein 126 is a novel factor involved in the negative regulation of DNA Damage response which is important for sustaining genomic integrity. | 28 |
| **SENP3** | SUMO specific peptidase 3. Redox sensor. Redox sensor that, when redistributed into nucleoplasm, that regulates HIF-1 transcriptional activity under oxidative stress. | 29,30 |
| **ANAPC11** | Anaphase promoting complex subunit 11. Subunit of the anaphase-promoting complex (APC). APC is activated during mitosis, remains active through most of G1, and is rapidly inactivated at the G1/S transition. | 31 |
| **PLBD2** | Phospholipase B domain containing 2. Involved in lipid catabolic process. Lysosomal localization | - |
| **CCNL2** | Cyclin L2.Regulator of the pre-mRNA splicing process, as well as in inducing apoptosis by modulating the expression of apoptotic and antiapoptotic proteins. Gene ID: 81669 | - |
| **PUM1** | Pumilio RNA binding family member 1.Participate in osteoporosis pathology and obesity | 32 |
| **ITPKB** | Inositol-trisphosphate 3-kinase B. Alzheimer's disease. Its overexpression is associated with increased cell death, enhanced astrogliosis, production of amyloid-β peptides and amyloid plaque formation. | 33 |
| **NAPA** | N-ethylmaleimide-sensitive factor (NSF) attachment protein alpha. Component of intracellular vesicle trafficking ensuring the continuity of vesicle fusion. It is implicated in regulation of cell survival because its overexpression protected cells from apoptosis induced by cytotoxic drugs | 34 |
| **IL15RA** | Interleukin 15 receptor subunit alpha. Atherosclerosis. Plasma Levels of Soluble Interleukin-2 Receptor α: Associations With Clinical Cardiovascular Events and Genome-Wide Association Scan | 35 |
| **ACSL1** | Acyl-CoA synthetase long chain family member 1. Key role in lipid biosynthesis and fatty acid degradation. Associated to fasting glucose and DM2 and also significantly associated with subclinical atherosclerosis. | 36–38 |
| **JMY** | Junction mediating and regulatory protein, p53 cofactor.P53 cofactor and controls actin dynamics in motile cells. It can affect apoptosis during the DNA damage response. Modulator of neuritogenesis. | 39–41 |
| **PUF60** | Poly(U) binding splicing factor 60.Loss-of-function variants cause a phenotype comprising growth/developmental delay and craniofacial, cardiac, renal, ocular and spinal anomalies, adding to disorders of human development resulting from aberrant RNA processing/spliceosomal function. Mutated in several cancers. | 42,43 |
| **CHSY1** | Chondroitin sulfate synthase 1. Involved in cell proliferation and morphogenesis. It may play a role in colorectal cancer, and mutations in this gene are a cause of temtamy preaxial brachydactyly syndrome. | 44 |
| **BAMBI** | BMP and activin membrane bound inhibitor. Obesity, fasting glucose changes over time. Diseases associated Wolfram Syndrome and Gnathodiaphyseal Dysplasia | 45,46 |

**Supplemental Table S5.** Functional analysis of the 21 loci, containing the 22 CpGs, significantly associated to IS using Fuma.

See Supplemental Tables, Excel file

**Supplemental Table S6.** GWAS association of the 21 loci, containing the 22 CpGs, significantly associated to IS. PheGenI information. Trait Related, single nucleotide polymorphism (SNP) associated to the trait; context, location in the gene; Gene and Gene ID associated; location, genomic position; P-Value associated to SNP; Source, source of information; pubmed, pubmed number where the results are published.

See Supplemental Tables, Excel file

**Supplemental Table S7. Characteristics of the 22 CpGs validated.** CpG id; Chr, chromosome location; pos, genomic position; gene, associated gene GeneHancer identifier, GeneHancer is a database of genome-wide enhancer-to-gene and promoter-to-gene associations, embedded in GeneCards: H3K27AC, histone modification associated to active gene expression: Dnase I, DNase I hypersensitive site (DHS), functionally related to transcriptional activity.

| **CpG** | **Chr** | **Pos (hg37)** | **CpG location** | **Gene** | **GeneHancer identifier** | **H3K27AC** | **DNAse I** |
|---|---|---|---|---|---|---|---|
| cg09915769 | 19 | 7660977 | Island | CAMSAP3 | GH19J007595 | no | yes |
| cg02463426 | 19 | 16683387 | Island | SLC35E1 | GH19J016568 | no | yes |
| cg00614832 | 16 | 73092394 | Island | ZFHX3 | GH16J073045 | no | yes |
| cg23962478 | 22 | 50354086 | Island | PIM3 | GH22J049957 | yes | yes |
| cg23681311 | 22 | 22221878 | Island | MAPK1 | GH22J021864 | yes | yes |
| cg13696351 | 9 | 140063617 | Island | LRRC26 | GH09J137166 | no | yes |
| cg01182555 | 14 | 62162064 | Island | HIF1A | no | no | yes |
| cg07691609 | 19 | 662740 | Island | RNF126 | GH19J000657 | yes | yes |
| cg01733795 | 17 | 7465439 | Island | SENP3 | GH17J007556 | yes | yes |
| cg08184047 | 17 | 79849980 | Island | ANAPC11 | GH17J081888 | yes | yes |
| cg04355250 | 12 | 113796401 | Island | PLBD2 | GH12J113356 | no | no |
| cg16573386 | 1 | 1334508 | Island | CCNL2 | GH01J001397 | yes | yes |
| cg23281075 | 1 | 31538727 | Island | PUM1 | GH01J031062 | yes | yes |
| cg07786668 | 16 | 73092391 | Island | ZFHX3 | GH16J073045 | no | yes |
| cg04482794 | 1 | 226925181 | Island | ITPKB | GH01J226735 | yes | yes |
| cg07806715 | 19 | 48018254 | Island | NAPA | GH19J047509 | no | yes |
| cg08676905 | 10 | 6019609 | Island | IL15RA | GH10J005974 | yes | yes |
| cg07619799 | 4 | 185747409 | Island | ACSL1 | GH04J184824 | yes | yes |
| cg04759220 | 5 | 78532560 | Island | JMY | GH05J079234 | yes | yes |
| cg01963056 | 8 | 144911482 | Island | PUF60 | GH08J143826 | yes | yes |
| cg25869317 | 15 | 101792241 | Island | CHSY1 | no | no | yes |
| cg04192862 | 10 | 28966472 | Island | BAMBI | GH10J028675 | no | yes |

**Supplemental Table S8. Meta-analysis.** CpG sites differentially methylated in relation to ischemic stroke. CpG id; Chr, chromosome location; genomic position; associated gene and observed mean β-values (standard deviation).

See Supplemental Tables, Excel file

**Supplemental Table S9.** List of genes harboring the 384 CpG associated to IS in the meta-analysis.

See Supplemental Tables, Excel file

**Supplemental Table S10.** Functional analysis of the 384 CpG associated to IS in the meta-analysis using Fuma.

See Supplemental Tables, Excel file

**Supplemental Table S11.** Association results of the meta-analysis loci. PheGenI information. Trait Related, single nucleotide polymorphism associated to the trait; context, location in the gene; Gene and Gene ID associated; location, genomic position; P-Value associated toSNP; Source, source of information; pubmed, pubmed number where the results are published.

See Supplemental Tables, Excel file

**Supplemental Table S12. Analysis by TOAST stratification.** Descriptive characteristics of the MAR_1 and MAR_2 samples stratified by TOAST. LAA, Large-Artery Atherosclerosis: CE, Cardiembolic; SVD, Small vessel diseases; UND, Undetermined IS subtypes; BMI, Body Mass Index; CHD, Coronary Heart Disease; NIHSS, stroke severity.

| **MAR samples** | Controls REGICOR | LAA | SVD | CE | Und | P-Value |
|---|---|---|---|---|---|---|
| N= 627 | N=224 | N=97 | N=94 | N=148 | N=64 | |
| Age, years* | 63 (57-70) | 70 (61-78) | 69 (54-77) | 79 (71-85) | 74 (67-82) | <0.001 |
| Gender, female, n (%) | 115 (50.4) | 21 (21) | 40 (42.1) | 89 (58.2) | 19 (29.2) | <0.001 |
| Hypertension, n (%) | 127 (55.7) | 75 (75) | 69 (72.6) | 126 (82.4) | 45 (69.2) | <0.001 |
| Smoking habit, n (%) | 92 (22) | 45 (45.9) | 31 (33) | 19 (12.5) | 17 (26.2) | <0.001 |
| BMI, kg/m2 * | 28.3 (25.8-31.6) | 26.6 (23.9-29.4) | 27 (24.1-30.9) | 26.8 (24.1-29.8) | 26.6(24-30.1) | 0.003 |
| Diabetes mellitus, n (%) | 40 (17.5) | 45 (45) | 34 (35.8) | 53 (34.9) | 24 (36.9) | <0.001 |
| Hyperlipidemia, n (%) | 100 (44.1) | 57 (57.6) | 50 (52.6) | 67 (43.8) | 42 (64.6) | 0.009 |
| Atrial fibrillation, n (%) | 8 (3.5) | 0 | 0 | 134 (87.6) | 15 (23.1) | <0.001 |
| CHD, n (%) | 0 | 9 (9) | 10 (10.5) | 29 (19.1) | 10 (15.4) | <0.001 |
| NIHSS† | - | 4 (2-10) | 3 (2-5) | 6(3.5-16.5) | 2 (0-5) | <0.001 |

**Supplemental Table S13. Atherotrombotic Stroke vs Controls analysis.** CpG sites differentially methylated in relation to ischemic stroke. CpG id; Chr, chromosome location; genomic position; associated gene and observed mean β-values (standard deviation).

See Supplemental Tables, Excel file

**Supplemental Table S14. Cardioembolic Stroke vs Controls analysis.** CpG sites differentially methylated in relation to ischemic stroke. CpG id; Chr, chromosome location; genomic position; associated gene and observed mean β-values (standard deviation).

See Supplemental Tables, Excel file

**Supplemental Table S15. Small vessel disease Stroke vs Controls analysis** CpG sites differentially methylated in relation to ischemic stroke. CpG id; Chr, chromosome location; genomic position; associated gene and observed mean β-values (standard deviation).

See Supplemental Tables, Excel file

**Supplemental Table S16. Undetermined Stroke vs Controls analysis** CpG sites differentially methylated in relation to ischemic stroke. CpG id; Chr, chromosome location; genomic position; associated gene and observed mean β-values (standard deviation).

See Supplemental Tables, Excel file

**Supplemental Table S17**. Functional analysis of the large-artery atherosclerosis (LAA) etiology subanalysis using Fuma.

See Supplemental Tables, Excel file

**Supplemental Table S18**. Functional analysis of the cardioembolic (CE) etiology subanalysis using Fuma.

See Supplemental Tables, Excel file

**Supplemental Table S19**. Functional analysis of the small vessel disease (SVD) etiology subanalysis using Fuma.

See Supplemental Tables, Excel file

**Supplemental Table S20**. Functional analysis of the undetermined (UND) etiology subanalysis using Fuma.

See Supplemental Tables, Excel file

**Supplemental Table S21.** GWAS association of the large-artery atherosclerosis etiology (LAA) subanalysis. PheGenI information. Trait Related, single nucleotide polymorphism (SNP) associated to the trait; context, location in the gene; Gene and Gene ID associated; location, genomic position; P-Value associated to SNP; Source, source of information; pubmed, pubmed number where the results are published.

See Supplemental Tables, Excel file

**Supplemental Table S22.** GWAS association of the cardioembolic (CE) etiology subanalysis subanalysis. PheGenI information. Trait Related, single nucleotide polymorphism (SNP) associated to the trait; context, location in the gene; Gene and Gene ID associated; location, genomic position; P-Value associated to SNP; Source, source of information; pubmed, pubmed number where the results are published.

See Supplemental Tables, Excel file

**Supplemental Table S23.** GWAS association of the small vessel disease (SVD) etiology subanalysis. PheGenI information. Trait Related, single nucleotide polymorphism (SNP) associated to the trait; context, location in the gene; Gene and Gene ID associated; location, genomic position; P-Value associated to SNP; Source, source of information; pubmed, pubmed number where the results are published.

See Supplemental Tables, Excel file

**Supplemental Table S24.** GWAS association of the undetermined (UND) etiology subanalysis. PheGenI information. Trait Related, single nucleotide polymorphism (SNP) associated to the trait; context, location in the gene; Gene and Gene ID associated; location, genomic position; P-Value associated to SNP; Source, source of information; pubmed, pubmed number where the results are published.

See Supplemental Tables, Excel file

12. R Core Team. R: A language and environment for statistical computing. *R*

*Foundation for Statistical Computing, Vienna, Austria*. 2014. Available at http://www.r-project.org/.

24. Eyileten C, Wicik Z, De Rosa S, Mirowska-Guzel D, Soplinska A, Indolfi C, Jastrzebska-Kurkowska I, Czlonkowska A, Postula M. MicroRNAs as Diagnostic and Prognostic Biomarkers in Ischemic Stroke—A Comprehensive Review and

Bioinformatic Analysis. *Cells*. 2018;7:249.

36. Draisma HHM, Pool R, Kobl M, et al. Genome-wide association study identifies

novel genetic variants contributing to variation in blood metabolite levels. *Nature communications*. 2015;6:7208.
